# Albumin-STING Nanoagonist Reprograms HSPCs to Antitumor Neutrophils Enhancing MHC I–Mediated CD8⁺ T Cell Immunity

**DOI:** 10.1101/2025.09.02.673154

**Authors:** Jinsong Tao, Hongyi Zhao, Chengyi Li, Hanning Wen, Fang Ke, Qiuxia Li, Miao He, Bo Wen, Zhongwei Liu, Wei Gao, Duxin Sun

**Affiliations:** Department of Pharmaceutical Sciences, College of Pharmacy, Rogel Cancer Center, University of Michigan, Ann Arbor, MI 48109, USA; Department of Pharmacology and Pharmaceutical Sciences, College of Pharmacy, University of Houston, TX, 77204, USA

## Abstract

Tumor-associated immunosuppressive neutrophils, termed myeloid-derived suppressor cells (MDSCs), compromise cancer immunotherapy. They can be pathologically programmed as early the hematopoietic stem and progenitor cell (HSPC) stage by suppressing interferon signaling. Reprogramming HSPCs toward antitumor neutrophils through the stimulator of interferon genes (STING) activation offers a promising therapeutic strategy. Here, we demonstrate that an albumin-STING nanoagonist (Nano ZSA-51D) reprograms HSPCs to generate antitumor neutrophils, enhancing MHC I-mediated CD8⁺ T cell immunity. Nano ZSA-51D activates STING–interferon signaling in HSPCs, promoting their expansion and differentiation toward granulocyte-monocyte progenitors via STING-NF-κB-IL-6 signaling. It further reprograms neutrophils into CD14⁺ICAM-1^+^ subset through STING-NF-κB–TNF-α signaling, enhancing tumor infiltration. These neutrophils upregulate interferon signaling and MHC I antigen presentation, boosting tumor-specific CD8⁺ T cell responses. Adoptive transfer of Nano ZSA-51D-reprogrammed neutrophils with α-PD1 therapy achieves complete colon tumor remission. Our findings provide a novel strategy to reprogram HSPCs toward antitumor neutrophils and highlight the potential of early interventions at HSPC stage to rewire neutrophil fate for cancer immunotherapy.

## INTRODUCTION

Tumor-associated immunosuppressive neutrophils, often referred to as myeloid-derived suppressor cells (MDSCs), are abundant in solid tumors that significantly compromise the efficacy of cancer immunotherapy in patients.^1,2^ This heterogeneous population is pathologically programmed by cancer-derived signals for immunosuppression.^3,4^ However, neutrophils can be programmed into either antitumor (N1) or protumor (N2) subsets.^5–7^ For instance, high neutrophil infiltration in tumors prior to immunotherapy is associated with poor prognosis, indicating an immunosuppressive role of neutrophils.^8,9^ While neutrophil infiltration induced by immunotherapy is correlated with improved therapeutic outcomes, suggesting an antitumor function of neutrophils.^10–19^ These antitumor neutrophils can directly kill tumor cells or act as antigen-presenting cells (APCs) to active T cell responses, thereby sensitizing tumors to anti-PD1 (α-PD1) therapy.^16–19^

Previous approaches have primarily focused on reprogramming neutrophils within the tumor microenvironment (TME) to reverse their immunosuppressive functions. Recent studies discover that the antitumor (N1) or protumor (N2) neutrophils may be programmed as early as the hematopoietic stem and progenitor cell (HSPC) stage, rather than solely through interconversion within the TME, due to the short lifespan of mature neutrophils.^20–25^ For example, tumor-derived signals can imprint immunosuppressive programming onto HSPCs, leading to the development of pro-tumor neutrophils.^20–22^ Conversely, certain immune interventions using Mycobacterium bovis Bacillus Calmette-Guérin (BCG) have been shown to reprogram HSPCs toward antitumor myeloid cells, including neutrophils and monocytes, that overcome tumor-induced immunosuppression.^23^ These findings highlight the therapeutic potential of reprogramming HSPCs toward antitumor neutrophils before exposure to cancer-derived signals.

Emerging evidence suggests that cancer-derived signals suppress the interferon response in granulocyte-monocyte progenitors (GMPs), reprogramming their differentiation toward immunosuppressive MDSC.^25^ The downregulation of interferon signaling is a key driver of their immunosuppressive phenotype.^26,27^ In contrast, interferon-stimulated neutrophils exhibit potent antitumor activity to sensitize α-PD1 immunotherapy in both mice and humans.^10,16^ Therefore, we aim to activate the stimulator of interferon genes (STING) pathway to restore the interferon signaling in both HSPCs and MDSCs. The STING pathway activation triggers phosphorylation of TBK1–IRF3 and NF-κB to induce interferons and proinflammatory cytokines such as TNF-a and IL-6, thereby promoting immunity.^28–30^ Notably, STING expression is higher in HSPCs than in their differentiated immune progeny, suggesting a unique function in early lineage programming.^31,32^ We hypothesize that STING agonist could activate interferon signaling in HSPCs and reprogram them toward antitumor neutrophils, thereby reversing tumor-driven MDSC development. Moreover, how STING activation reprograms HSPCs to antitumor neutrophils and the underlying antitumor mechanisms of these neutrophils for cancer immunotherapy remain elucidated.

Here, we developed an albumin-STING nanoagonist (Nano ZSA-51D) that reprograms HSPCs to antitumor neutrophils with enhanced MHC I antigen presentation, which migrate from bone marrow into tumors to remodel the TME and promote CD8⁺ T cell immunity (**Fig. 1a**). The dimeric STING agonist ZSA-51D was synthesized from our monomeric STING agonist ZSA-51^33^ and encapsulated into mouse albumin to generate Nano ZSA-51D. Nano ZSA-51D activated the STING–interferon signaling in HSPCs, promoting their expansion and skewing differentiation toward GMPs through STING-NF-κB-IL-6 signaling in bone marrow. It further reprogrammed immature (CD101^-^) and mature (CD101^+^) neutrophils into CD14⁺ICAM-1^+^ subsets through STING–NF-κB–TNF-α signaling, thereby enhancing tumor infiltration. Transcriptomic and immune profiling revealed that these neutrophils upregulated interferon signaling and MHC I antigen presentation to boost tumor-specific CD8^+^ T cell responses. Importantly, adoptive transfer of Nano ZSA-51D-reprogrammed neutrophils in combination with α-PD1 therapy achieved complete remission of colon tumors with durable immune memory. Nano ZSA-51D combined with α-PD1 exhibited potent efficacy in colon cancer and pancreatic cancer models.

**Figure 1.**
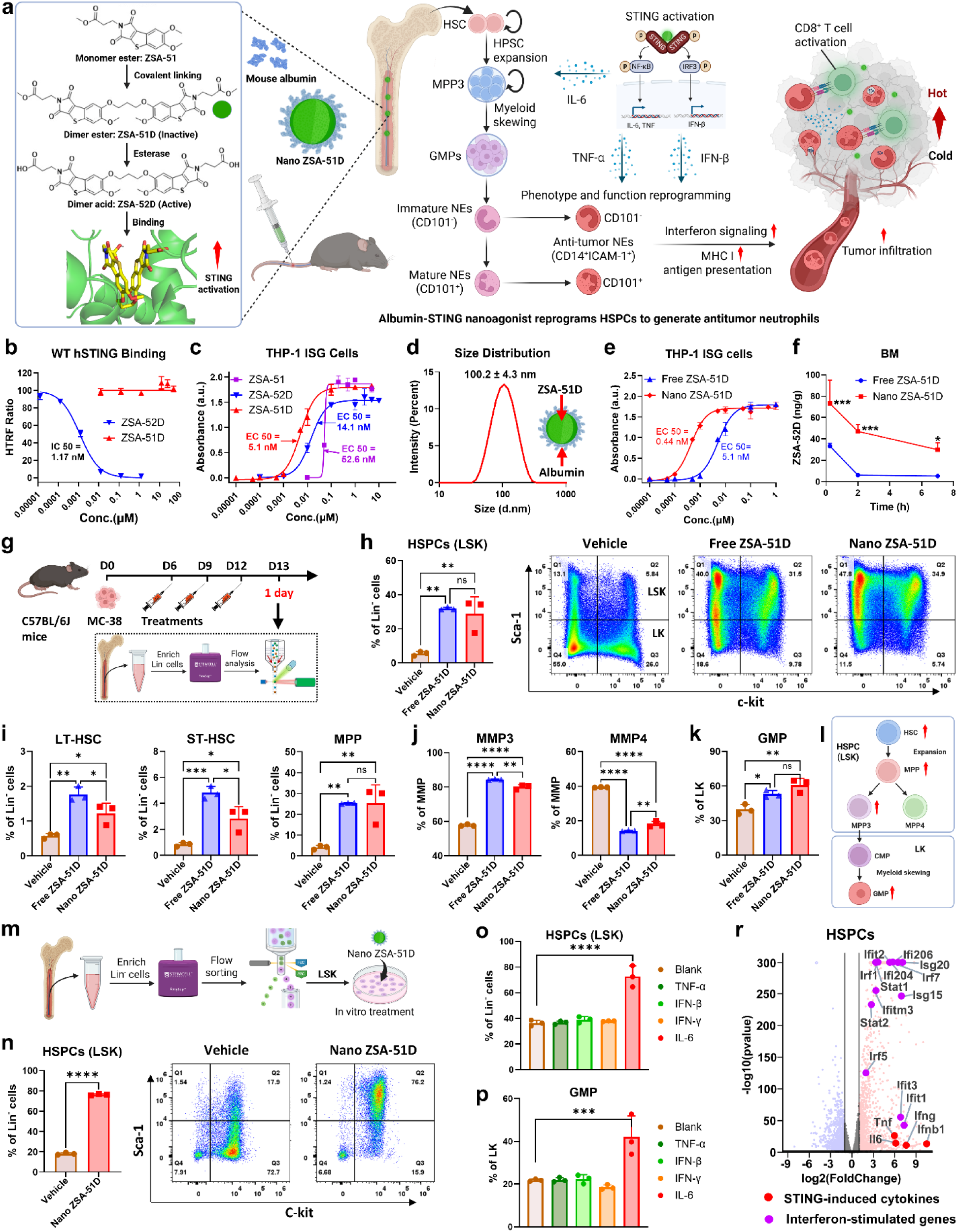
Nano ZSA-51D potently activates STING pathway to expand HSPCs and promote GMP differentiation with strong interferon signaling. **(a)** Schematic illustration of Nano ZSA-51D that reprograms HSPCs into antitumor neutrophils for cancer immunotherapy. Dimeric STING agonist ZSA-51D derived from monomeric STING agonist ZSA-51 was encapsulated into mouse albumin to form Nano ZSA-51D. Chemical structures of ZSA-51, ZSA-51D and its acid form, ZSA-52D, along with molecular docking simulation showing ZSA-52D (yellow) binding to the STING protein (green, PDB: 6UKZ), performed using AutoDock. Nano ZSA-51D expands HSPCs and skews GMP differentiation via STING–NF-κB–IL-6 signaling, which further reprograms immature (CD101⁻) and mature (CD101⁺) bone marrow neutrophils into CD14⁺ICAM-1⁺ subsets through STING–NF-κB–TNF-α signaling, upregulating interferon signaling and MHC I antigen presentation to promote CD8⁺ T cell antitumor responses. **(b)** Binding affinity of ZSA-51D and ZSA-52D to wild-type (WT) human STING (hSTING) protein, determined by Homogeneous Time-Resolved Fluorescence (HTRF) assay (n=3). **(c)** STING pathway activation was assessed in THP1-Blue ISG reporter cells with virous concentrations of ZSA-51D and ZSA-52D for 24 hours, demonstrating induction of the STING– IRF signaling cascade (n=3). **(d)** Particle size distribution and schematic structure of Nano ZSA-51D. **(e)** Comparison of STING-IRF pathway activation by free and Nano ZSA-51D in THP1-Blue ISG reporter cells after 24-hour treatment (n=3). **(f)** Pharmacokinetic profiles of Free and Nano ZSA-51D (1 mg/kg, I.V.), showing concentrations of ZSA-52D (acid form) in bone marrow at 15 minutes, 2 hours and 7 hours post-injection, quantified via LC-MS/MS (n = 3). * p < 0.05, *** p < 0.001 vs. Free ZSA-51D. **(g)** Schematic illustration of treatment schedule and flow cytometry analysis to examine the composition of HSPC subsets in bone marrow of MC-38 tumor bearing mice 24 hours post-treatment. The flow cytometry analysis of HSPCs was performed after enrichment of Lin^-^ cells. **(h)** Quantification (left) and representative flow cytometry plots (right) of HSPCs (Lin^-^Sca-1^+^c-Kit^+^, LSK) within Lin^-^ cells in bone marrow following free or Nano ZSA-51D treatment (n = 3). **p < 0.01, ns: not significant. **(i)** Quantification (left) of LT-HSC (CD150^+^CD48^-^), ST-HSC (CD150^-^CD48^-^) and MPP (CD150^-/+^CD48^+^) within Lin^-^ cells in bone marrow following free or Nano ZSA-51D treatment (n = 3). *p < 0.05, **p < 0.01, ***p < 0.001, ns: not significant. **(j)** Quantification of myeloid-biased multipotent progenitor 3 (MPP3, CD34^+^CD135^-^) and lymphoid-biased multipotent progenitor 4 (MPP4, CD34^+^CD135^+^) within MMP (Lin^-^Sca-1^+^c-Kit^+^CD150^-/+^CD48^+^) in bone marrow following free or Nano ZSA-51D treatment (n = 3). ****p < 0.0001, ns: not significant. **(k)** Quantification of granulocyte-monocyte progenitor (GMP, CD150^-^CD16/32^+^) within LK (Lin^-^ Sca-1^-^c-Kit^+^) in bone marrow following free or Nano ZSA-51D treatment (n = 3). *p < 0.05, **p < 0.01, ns: not significant. **(l)** Schematic illustration of HSPCs expansion and skewing differentiation toward GMP after STING agonist treatments. HSC, hematopoietic stem cell; MPP, multipotent progenitor; MPP3, myeloid-biased multipotent progenitor 3; MPP4, lymphoid-biased multipotent progenitor 4; CMP, common myeloid progenitor; GMP (granulocyte-monocyte progenitor). **(m)** Experimental schematic of the in vitro treatment of LSK via Nano ZSA-51D (100 nM). The LSK (Lin^-^Sca-1^+^c-Kit^+^) was sorted by flow after enrichment of Lin-cells from bone marrow cells. **(n)** Quantification (left) and representative flow cytometry plots (right) of LSK (Lin^-^Sca-1^+^c-Kit^+^) within Lin^-^ cells following in vitro treatment of Nano ZSA-51D for 3 days (n = 3). ****p < 0.0001. **(o, p)** Quantification (left) and representative flow cytometry plots (right) of HSPCs (Lin^-^Sca-1^+^c-Kit^+^, LSK) within Lin^-^ cells (i) and GMP (CD150^-^CD16/32^+^) within LK (Lin^-^Sca-1^-^c-Kit^+^) (j) following in vitro treatments of the STING pathway downstream cytokines (TNF-α, IFN-β, IFN-γ and IL-6) for 3 days (n = 3). ***p < 0.001, ****p < 0.0001. **(r)** Volcano plot of differentially expressed gene analysis between vehicle and Nano ZSA-51D treated HSPCs. Significance threshold for P adjusted value was 0.01 and foldchange was 2. The STING-induced cytokines (red dots) and interferon-stimulated genes (purple dots) were highlighted.

## RESULTS

### Albumin-STING Nanoagonist (Nano ZSA-51D) Potently Activates STING Signaling Pathway and Enhances Bone Marrow Delivery

Building on our previously developed monomeric STING agonist, ZSA-51,^33^ we synthesized a novel non-CDN dimeric STING agonist, ZSA-51D, which exhibited superior STING pathway activation (**Fig. 1a, Left**). ZSA-51D is an ester-based prodrug, that requires hydrolysis by esterases to convert into its active acid form (ZSA-52D) before binding to STING protein. Both ZSA-51D and ZSA-52D were synthesized (**Fig. S1)**. Due to its enhanced cell permeability, ZSA-51D can be easily uptake inside cells and efficiently converted to ZSA-52D, resulting in potent and sustained STING activation. As anticipated, ZSA-51D showed no binding with human STING, whereas the activated form ZSA-52D displayed a potent binding affinity to human STING, with an IC_50_ of 1.17 nM (**Fig. 1b**). In THP1-Blue™ ISG reporter cells, ZSA-51D demonstrated a superior EC_50_ of 5.1 nM, outperforming ZSA-52D (EC_50_ = 14.1 nM) and monomer ZSA-51 (EC_50_ = 52.6 nM) (**Fig. 1c**).

To facilitate systemically intravenous administration, we encapsulated ZSA-51D into an albumin nanoparticle (Nano ZSA-51D).^34–36^ The average particle size of Nano ZSA-51D was 100.2 ± 4.3 nm (**Fig. 1d**). Notably, Nano ZSA-51D exhibited an 11.6-fold enhancement in STING activation (EC_50_ = 0.44 nM) compared to free ZSA-51D (EC_50_ = 5.1 nM) in THP1-Blue^TM^ ISG cells (**Fig. 1e**). The pharmacokinetics studies showed that Nano ZSA-51D enhanced bone marrow accumulation compared to free ZSA-51D (**Fig. 1f**), whereas drug distribution in tumor and systemic circulation was similar between the two groups (**Fig. S2**). Western blot analysis further confirmed that Nano ZSA-51D significantly enhanced phosphorylation of STING and downstream effectors IRF3 and NF-κB p65 in bone marrow cells, thereby promoting the production of IFN-β, TNF-α and IL-6 (**Fig. S3**).^30^ These results indicate that Nano ZSA-51D potently activates both the STING-IRF3 and STING-NF-κB pathways.

### Nano ZSA-51D Expands HSPCs and Skews Differentiation Toward GMPs

Neutrophils are derived from HSPCs in the bone marrow through a series of progressively restrictive differentiation steps (**Fig. S4A**). To further explore whether Nano ZSA-51D–mediated STING activation effects on HSPCs in vivo, we analyzed the composition of HSPC subsets in bone marrow of MC-38 colon tumor-bearing mice 24 hours after 3 doses (1 mg/kg, I.V.) (**Fig. 1g**).

Surprisingly, both free and Nano ZSA-51D treatments resulted in a marked expansion of the HSPCs (Lin^-^Sca-1^+^c-Kit^+^, LSK) (**Fig. 1h**), which includes long-term hematopoietic stem cell (LT-HSC), short-term HSC (ST-HSC) and multipotent progenitor (MPP) (**Fig. 1i and Fig. S4B**).^37^ Notably, MPP exhibited the most dramatic expansion, rising from 4.1% to 25.3%. Among the MPP, the myeloid-biased multipotent progenitor 3 (MPP3) highly increased, while the lymphoid-biased multipotent progenitor 4 (MPP4) decreased, respectively (**Fig. 1j and Fig. S4C**). In downstream progenitors, we also observed an increase in granulocyte-monocyte progenitor (GMP) (**Fig. 1k and Fig. S4D**). These results suggest that Nano ZSA-51D expands HSPCs and skews differentiation toward granulocyte-monocyte progenitors (GMP), thereby potentially enhancing myeloid lineage output and supporting innate immune responses (**Fig. 1l**).

### Nano ZSA-51D Expands HSPCs and Skews Differentiation Toward GMP via STING-NF-κB-IL-6 Signaling

To elucidate how STING agonists can expand HSPCs and skew differentiation toward GMP, we isolated the HSPCs (LSK cells) from the bone marrow of MC-38 tumor-bearing C57BL/6J mice and treated them with Nano ZSA-51D (100 nM) for 3 days in vitro (**Fig. 1m**). Interestingly, Nano ZSA-51D dramatically expanded HSPCs (LSK populations) from 18.1% to 76.3% (**Fig. 1n**), while vehicle-treated groups exhibited the expected differentiation of HSPCs from LSK to LK populations. This observation suggests that STING activation can directly expand HSPCs at stem and multipotent progenitor states.

Since STING activation can induce different downstream cytokines such as IFN-β, TNF-α, IFN-γ and IL-6, we further test which of these cytokines can expand the HSPCs and skew differentiation toward GMP. We treated the isolated HSPCs (LSK cells) with cytokines for 3 days and monitored the composition changes of HSPCs. Among these cytokines, only IL-6 significantly expanded HSPCs (LSK population) (**Fig. 1o and Fig. S5A)** and increased the GMP population (**Fig. 1p and Fig. S5B**). Given that IL-6 is a key downstream effector of the STING–NF-κB axis, these results suggest that Nano ZSA-51D initiates STING-NF-κB-IL-6 signaling to expand HSPCs, and skew differentiation toward the GMP.

### Nano ZSA-51D Activates STING–Interferon Signaling in HSPCs

Cancer-derived signals can reprogram GMPs derived from HSPCs to promote immunosuppressive differentiation of MDSCs by suppressing interferon response.^25–27^ To investigate whether Nano ZSA-51D activates the STING pathway to upregulate the interferon signaling in HSPCs, we performed mRNA sequencing on vehicle- and Nano ZSA-51D-treated HSPCs. Nano ZSA-51D significantly increased the expression of STING-induced cytokines, including IFN-β, TNF-α, IFN-γ and IL-6, and upregulated multiple interferon-stimulated genes, such as Irf1, Irf5, Irf7, Stat1, Stat2, Ifit1, and Isg15, among others (**Fig. 1r**). These results indicate that Nano ZSA-51D robustly activates the STING–interferon signaling in HSPCs, thereby reprogramming their transcriptional landscape toward an antitumor immune phenotype.

### Nano ZSA-51D Promotes Neutrophil Generation and Tumor Infiltration

To determine whether Nano ZSA-51D–mediated HSPC expansion and GMP differentiation promote neutrophil generation and tumor infiltration, we analyzed the neutrophils in bone marrow, blood and tumors of MC-38 colon tumor-bearing mice 24 hours after 3 doses (1 mg/kg, I.V.) (**Fig. S6A**).

In the bone marrow, both free and Nano ZSA-51D significantly increased proportion of neutrophils among CD45^+^ cells from 50.1% to 59.7% and 63.3%, respectively (**Fig. S6B**). In peripheral blood, Nano ZSA-51D also elevated circulating neutrophils, but which was less than free ZSA-51D (**Fig. S6C**). Interestingly, in MC-38 tumors, Nano ZSA-51D dramatically increased tumor-infiltrating neutrophils from 11.2% to 56.7% of CD45^+^ immune cells, a substantially greater increasement than free ZSA-51D (**Fig. S6D**). Collectively, these results demonstrate that while both free and Nano ZSA-51D promote granulopoiesis to generate neutrophils, Nano ZSA-51D more effectively drives their infiltration into tumors.

### Nano ZSA-51D Reprograms Bone Marrow Neutrophils into CD14⁺ICAM-1^+^ Subset to Enhance Tumor Infiltration

Recent studies have demonstrated that the immature/activated (CD101^-^CD14^+^) subset of tumor-infiltrating neutrophils is associated with improved responses to immunotherapy.^10,38–40^ CD101 is the maturation marker of neutrophils and CD14 is the activation marker of pattern recognition receptor-mediated immune responses.^41–45^ To understand how Nano ZSA-51D further reprograms bone marrow neutrophils for tumor infiltration and antitumor activity, we analyzed neutrophil subtypes in the bone marrow, peripheral blood, and tumors of MC-38 tumor-bearing mice 24 hours following STING agonist treatment.

In the bone marrow of vehicle-treated mice, most neutrophils were immature/inactivated (CD101⁻CD14^-^). Nano ZSA-51D treatment lead a 4-fold increase in immature/activated (CD101^-^ CD14^+^) neutrophils from 7.5% to 29.6 %, which was higher than free ZSA-51D treatment (**Fig. 2a**).

**Figure 2.**
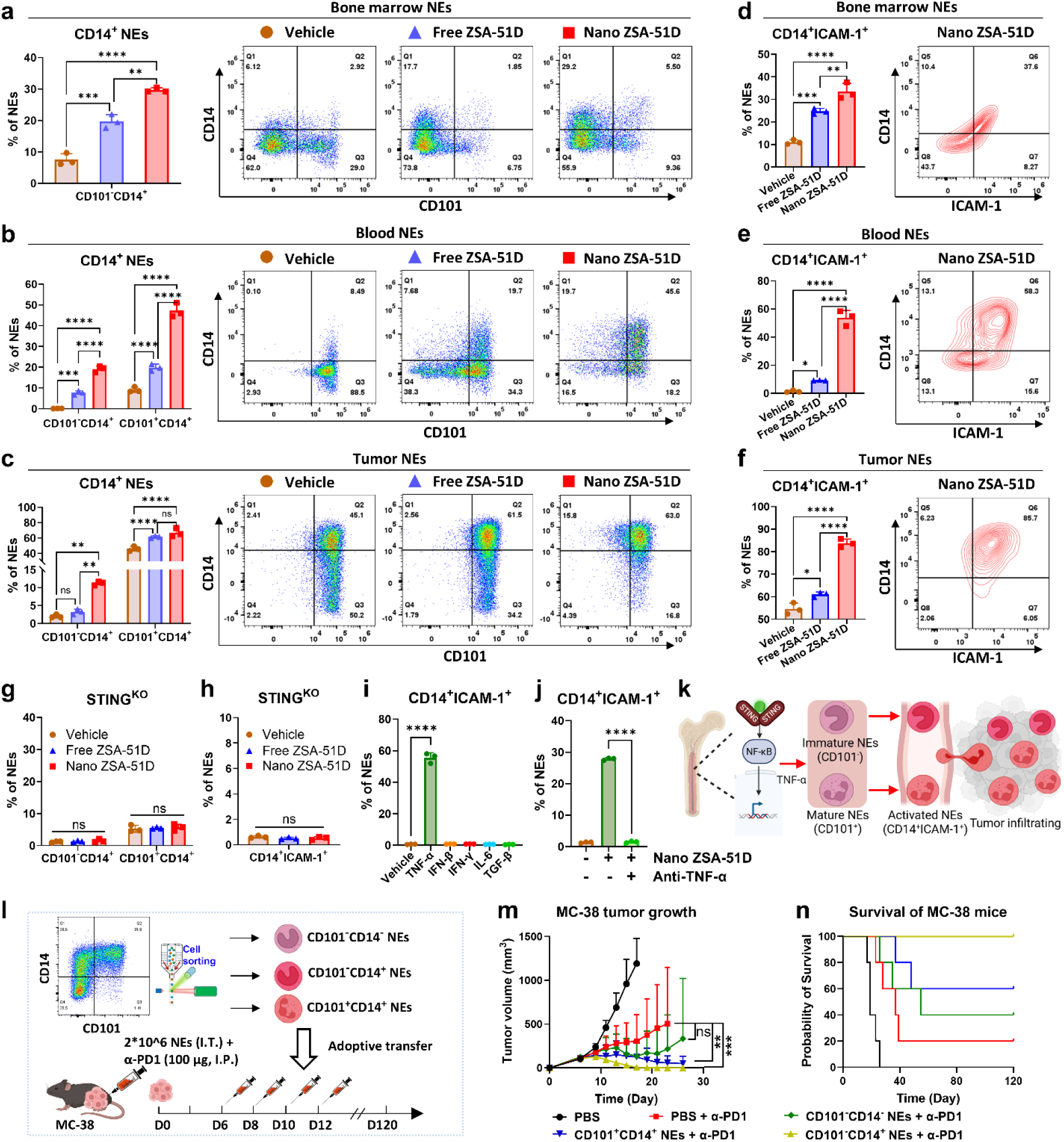
Nano ZSA-51D reprograms bone marrow neutrophils into CD14⁺ICAM-1^+^ antitumor subset via STING–NF-κB–TNF-α signaling, driving complete MC-38 tumor remission with α-PD1 therapy. **(a-c)** Quantification (left) and representative flow cytometry plots (right) of CD101^-^CD14^+^ and CD101^+^CD14^+^ neutrophil (NE) subtypes in bone marrow (a), blood (b) and tumors (c) of MC-38 tumor-bearing C57BL/6J mice 24 hours after systemic administration of free and Nano ZSA-51D (1 mg/kg) (n = 3). **p < 0.01, ***p < 0.001, ****p < 0.0001, ns: not significant. **(d-f)** Quantification (left) and representative flow cytometry plots (right) of CD14^+^ICAM-1^+^ neutrophil subtypes in bone marrow (d), blood (e) and tumors (f) after systemic administration of Nano ZSA-51D (1 mg/kg) (n = 3). *p < 0.05, **p < 0.01, ***p < 0.001, ****p < 0.0001. **(g-h)** Quantification of CD101^-^CD14^+^, CD101^+^CD14^+^ (g) and CD14^+^ICAM-1^+^ (h) neutrophils after overnight in vitro treatments with free and Nano ZSA-51D (100 nM) in bone marrow cells from STING knockout mice (C57BL/6J-Sting1gt/J) (n = 3). ns: not significant. **(i)** Quantification of CD14^+^ICAM-1^+^ neutrophils following in vitro treatments of the STING pathway downstream cytokines (TNF-α, IFN-β, IFN-γ and IL-6) for overnight. The immunosuppressive TGF-β as control (n = 3). ****p < 0.0001. **(j)** Quantification and changes of CD14⁺ICAM-1⁺ neutrophils after overnight in vitro treatment with Nano ZSA-51D (100 nM) and TNF-α neutralization (10 μg/ml anti-TNF-α) (n=3). ****p < 0.0001. **(k)** Schematic illustration of STING activation-induced reprogramming of immature (CD101^-^) and mature (CD101^+^) bone marrow neutrophils into activated CD14^+^ICAM-1^+^ subsets with enhanced tumor infiltration. **(l)** Schematic experiment of neutrophil adoptive transfer. CD101^-^CD14^-^, CD101^-^CD14^+^ and CD101^+^CD14^+^ neutrophils (NEs) were sorted from bone marrow cells pretreated overnight with Nano ZSA-51D (100 nM), then intratumorally injected into MC-38 tumor-bearing mice alongside intraperitoneal α-PD1 (100 μg/mouse). (**m, n**) Tumor growth curves (l) and survival curves (m) of MC-38 tumor-bearing mice after adoptive transfer of CD101^-^CD14^-^, CD101^-^CD14^+^ or CD101^+^CD14^+^ neutrophils in combination with α-PD1 (100 μg/mice) treatments (n=5). **p < 0.01, ***p < 0.001, ns: not significant. Survival comparisons were performed using a log-rank (Mantel–Cox) test.

In the blood circulation of vehicle-treated mice, most neutrophils were mature/inactivated (CD101^+^CD14^-^) (**Fig. 2b)**. While Nano ZSA-51D treatment dramatically increased the immature/activated (CD101^-^CD14^+^) neutrophils from 0.08% to 20%, and mature/activated (CD101^+^CD14^+^) neutrophils from 9.1% to 47.2%, which were both substantially higher than free ZSA-51D treatment. These data suggest that Nano ZSA-51D strongly activates both immature and mature CD14^+^ neutrophils, promoting their migration from bone marrow into bloodstream.

In the tumors, activated CD14⁺ neutrophils preferentially infiltrated tumors following Nano ZSA-51D treatment. Specifically, the proportion of immature/activated CD101⁻CD14⁺ neutrophils increased from 2.0% to 11.2%, and mature/activated CD101⁺CD14⁺ neutrophils rose from 46.1% to 66.8% (**Fig. 2c**). In contrast, although free ZSA-51D also expanded total neutrophil numbers in bone marrow and blood (**Fig. S6**), it induced fewer CD14⁺ subsets, resulting in less tumor infiltration. In addition, Nano ZSA-51D induced CD14^+^ neutrophils highly expressed intercellular adhesion molecule-1 (ICAM-1, CD54) in bone marrow and blood circulation, which promote CD14^+^ICAM-1^+^ neutrophil infiltration into tumor (**Fig. 2d-f** and **Fig. S7**).^46,47^ These data suggest that Nano ZSA-51D effectively reprograms immature (CD101^-^) and mature (CD101^+^) bone marrow neutrophils into activated CD14⁺ICAM-1^+^ subsets that enhance infiltration into tumors.

### Nano ZSA-51D Reprograms Neutrophils into CD14⁺ICAM-1^+^ Subset through STING–NF-κB–TNF-α Signaling

To determine whether Nano ZSA-51D inducing CD14^+^ICAM-1^+^ neutrophils is mediated by STING pathway activation, we isolated bone marrow cells from wild-type (WT) C57BL/6J and STING knockout (C57BL/6J-Sting1gt/J) mice and treated them with free or Nano ZSA-51D (100 nM) in vitro for overnight. As expected, Nano ZSA-51D (100 nM) significantly increased both CD101^-^CD14^+^ and CD101^+^CD14^+^ neutrophil populations with high ICAM-1 expression (CD14^+^ICAM-1^+^) from WT C57BL/6J mice, exceeding the effects observed with free ZSA-51D (**Fig. S8A, B**). However, bone marrow neutrophils from STING knockout mice failed to reprogram neutrophiles into CD101^-^CD14^+^, CD101^+^CD14^+^ and CD14^+^ICAM-1^+^ neutrophil populations, confirming the requirement of STING activation (**Fig. 2g, h and Fig. S8C, D**).

To further study which downstream mediators of STING signaling play a role in reprograming neutrophil subtypes, we treated the isolated bone marrow neutrophils with TNF-α, IFN-β, IFN-γ and IL-6. We also used TGF-β as a control for immunosuppressive signaling. Among these, only TNF-α effectively reprogrammed bone marrow neutrophils into CD101^-^CD14^+^, CD101^+^CD14^+^ and CD14^+^ICAM-1^+^ subsets (**Fig. 2i and Fig. S9A-D**). After TNF-α neutralization with antibody treatment, Nano ZSA-51D no longer reprogrammed neutrophils into the CD14⁺ICAM-1⁺ subset (**Fig. 2j and Fig. S9E**). Given that TNF-α is a key effector of STING-NF-κB axis, these findings suggest that Nano ZSA-51D reprograms immature and mature bone marrow neutrophils into CD14^+^ICAM-1^+^ subsets via STING-NF-κB-TNF-α signaling, enabling their infiltration into tumors (**Fig. 2k**).

### Nano ZSA-51D-Reprogrammed Neutrophils Drive Complete MC-38 Tumor Remission with α-PD1 Therapy

To determine whether Nano ZSA-51D-reprogrammed bone marrow neutrophils possess antitumor activity, we isolated three neutrophil subsets from bone marrow cells treated with Nano ZSA-51D (100 nM) in vitro: immature/inactivated (CD101⁻CD14⁻), immature/activated (CD101⁻CD14⁺) and mature/activated (CD101⁺CD14⁺). Each subset was adoptively transferred intratumorally (I.T.) into MC-38 tumors in combination with α-PD1 therapy (**Fig. 2l**).

Interestingly, adoptive transfer of immature/activated (CD101^-^CD14^+^) neutrophils combined with α-PD1 achieved 100% tumor complete remission (100% CR) without recurrence, resulting in 100% long-term survival (**Fig. 2m, n and Fig. S10**). The mature/activated (CD101^+^CD14^+^) neutrophils also significantly enhanced anti-tumor efficacy, achieving a 60% CR rate. In contrast, the immature/inactivated (CD101^-^CD14^-^) neutrophils with α-PD1 did not significantly improve therapeutic outcomes compared to α-PD1 monotherapy. These findings suggest that Nano ZSA-51D-reprogrammed CD14^+^ neutrophils possess potent antitumor properties that synergize with α-PD1 therapy.

To investigate whether the anti-tumor effect conferred by CD101⁻CD14⁺ neutrophils involved long-term immune memory, mice that achieved complete remission were rechallenged with MC-38 tumor cells 120 days post-treatment (**Fig. S11A**). Notably, these mice completely resisted tumor re-growth (**Fig. S11B**), indicating the establishment of durable antitumor immune memory.

To further characterize the immune memory response, we analyzed CD8⁺ and CD4⁺ T cell memory subsets in the spleen, lymph nodes and blood of cured MC-38 mice at 180-day post-rechallenge (**Fig. S11A**). Flow cytometry analysis revealed a significant increase in CD8⁺ T central memory (TCM) cells in cured MC-38 mice, with a 1.53-, 2.40- and 1.84-fold increase in spleen, lymph nodes and blood compared to naïve mice (**Fig. S11C, E, F**). CD4^+^ T effector memory (TEM) cells also increased in all tissues examined, while CD4^+^ TCM cells remained unchanged (**Fig. S11D, F, H**). These results demonstrate that adoptive transfer of CD101⁻CD14⁺ neutrophils with α-PD1 therapy promotes long-term CD8^+^ T cell memory, driving complete tumor remission and long-lasting tumor immunity.

### Nano ZSA-51D Upregulates Interferon Signaling and MHC I Antigen Presentation in Neutrophils

To further elucidate how Nano ZSA-51D-reprogrmmed neutrophils contribute to anticancer efficacy, we performed bulk mRNA sequencing on sorted total neutrophils and their subtypes from bone marrow cells following Nano ZSA-51D (100 nM) treatment in vitro. Differential gene expression analysis revealed that antitumor neutrophil (N1) markers were upregulated, while protumor (N2) markers were downregulated following treatment, indicating a shift toward an N1-like neutrophil phenotype^48–50^ (**Fig. 3a**).

**Figure 3.**
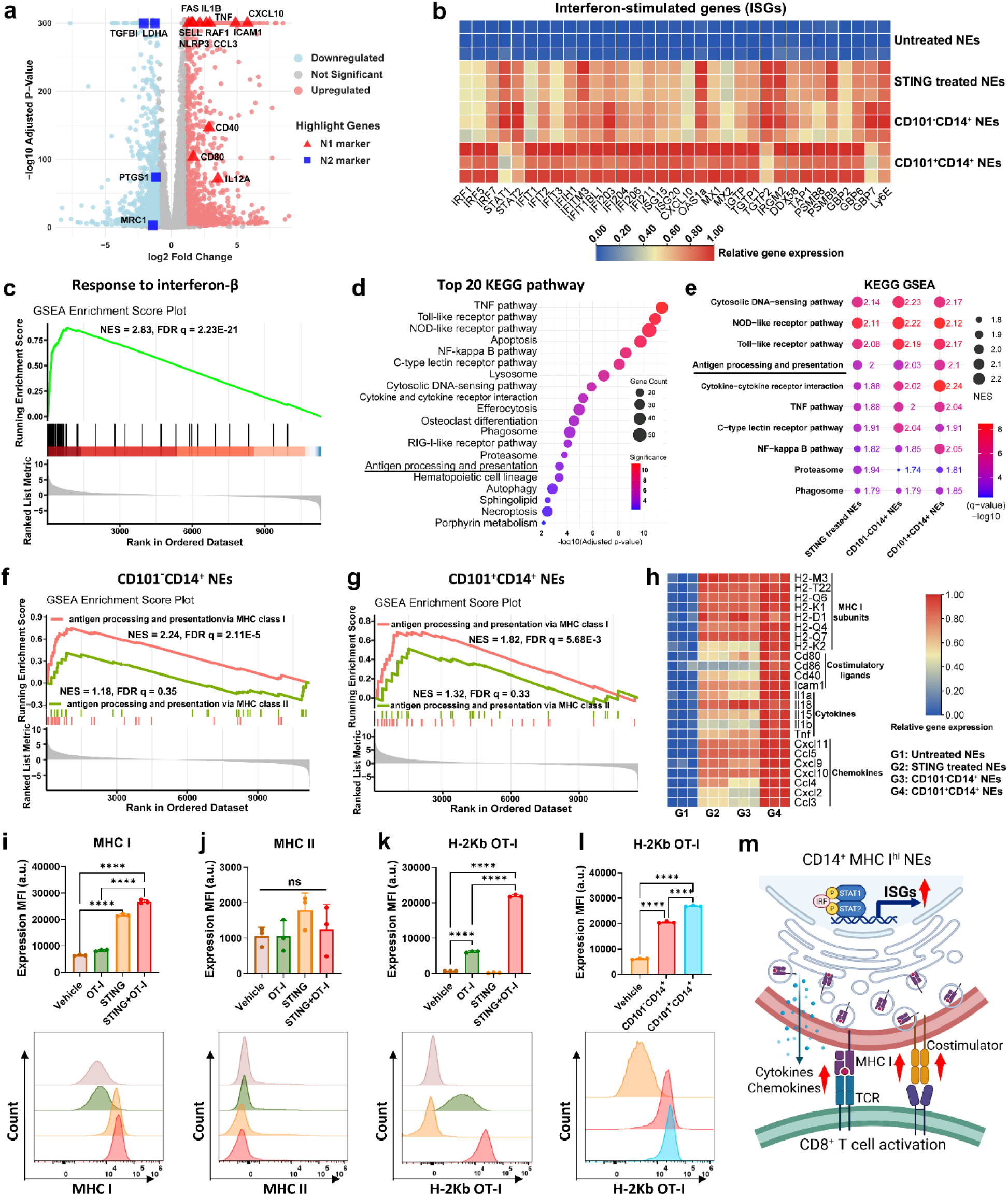
Nano ZSA-51D enhances interferon signaling and MHC I antigen presentation in neutrophils. **(a)** Volcano plot of differentially expressed gene analysis between vehicle and Nano ZSA-51D treated neutrophils. Significance threshold for P adjusted value was 0.01 and foldchange was 2. The antitumor (N1, red triangle) and protumor (N2, blue square) neutrophil markers were highlighted. **(b)** Heatmap of relative interferon-stimulated gene (ISG) expression in Nano ZSA-51D-treated neutrophils (NEs) and their CD101^-^CD14^+^ and CD101^+^CD14^+^ subtypes, compared to untreated neutrophils. **(c)** Gene Ontology (GO) GESA enrichment score plots for biological process that response to interferon-beta (IFN-β) in Nano ZSA-51D treated neutrophils. **(d)** Top 20 KEGG pathway enrichment analysis based on the differentially expressed genes (DEGs) in neutrophils from Nano ZSA-51D-treated and vehicle-treated groups, ranked by adjusted p-values. **(e)** Top 10 upregulated pathways in Nano ZSA-51D-treated neutrophils and their CD101^-^CD14^+^ and CD101^+^CD14^+^ subtypes, identified by KEGG Gene Set Enrichment Analysis (GESA). Normalized enrichment score (NES > 1.5) indicates significant upregulation. FDR q-value (q < 0.05) denotes statistical significance after false discovery rate adjustment. (**f, g**) Gene Ontology (GO) GESA enrichment score plots for antigen processing and presentation via MHC class I and II in CD101^-^CD14^+^ (f) and CD101^+^CD14^+^ (g) neutrophils. Normalized enrichment score (NES > 1.5) indicates significant upregulation. FDR q-value (q < 0.05) denotes statistical significance after false discovery rate adjustment. **(h)** Heatmap of relative gene expression of MHC class I subunits, costimulatory ligands, T cell-activating cytokines and T cell-recruiting chemokines in Nano ZSA-51D-treated neutrophils and their CD101^-^CD14^+^ and CD101^+^CD14^+^ subtypes. (**i, j**) Protein expression of MHC class I (i) and II (j) molecules on bone marrow neutrophils following treatments with OT-I or/and Nano ZSA-51D (n = 3). ****p < 0.0001, ns: not significant. (k) Presentation of OT-I peptide via MHC class I (H-2K^b^-OT-I complex) on neutrophils following treatments with OT-I or/and Nano ZSA-51D (n = 3). ****p < 0.0001. (l) Presentation of OT-I peptide via MHC class I (H-2K^b^-OT-I complex) on CD101^-^CD14^+^ and CD101^+^CD14^+^ neutrophils following OT-I treatment (n = 3). ****p < 0.0001. (m) Schematic illustration of antigen presentation by ISG-expressing CD14^+^MHC I^hi^ neutrophils via MHC class I to activate CD8^+^ T cells.

Moreover, the interferon-stimulated genes (ISGs) were highly upregulated in neutrophils following Nano ZSA-51D treatment (**Fig. 3b**). GO Gene set enrichment analysis (GSEA) further revealed significant enrichment of the gene response to IFN-β (NES = 2.83, FDR q = 2.23E-21) in Nano ZSA-51D treated neutrophils (**Fig. 3c**), indicating that STING pathway activation drives IFN-β–mediated interferon signaling in these cells.

KEGG pathway enrichment analysis identified the top 20 hallmark pathways altered in neutrophils by Nano ZSA-51D (**Fig. 3d**). Among them, the key innate immune signaling pathways were highly enriched and upregulated, including Cytosolic DNA-sensing, NOD-like receptor, toll-like receptor and C-type lectin receptor pathways, along with their downstream signaling cascades such as the TNF and NF-kappa B pathways (**Fig. 3e and Fig. S12A, B**). These pathways play critical roles in enabling the CD14^+^ neutrophils to detect pathogen-associated molecular patterns (PAMPs) and damage-associated molecular patterns (DAMPs) in the TME.^51–54^

A striking finding was the significant enrichment and upregulation of antigen processing and presentation pathway (NES = 2.00, FDR q = 4.36E-5) in neutrophils by Nano ZSA-51D treatment, as well as immature/activated (CD101^-^CD14^+^) (NES = 2.03, FDR q = 3.49E-5) and mature/activated (CD101^+^CD14^+^) (NES = 2.1, FDR q = 2.24E-6) neutrophils (**Fig. 3d, e and Fig. S12C)**. Further GO GSEA analysis confirmed that MHC class I-mediated antigen processing and presentation were highly upregulated in CD101^-^CD14^+^ and CD101^+^CD14^+^ neutrophils (**Fig. 3f, g)**. In contrast, MHC class II-mediated antigen processing and presentation remained unchanged (NES < 1.5, FDR q > 0.05).

Consistently, Nano ZSA-51D markedly upregulated the expression of MHC class I subunits, costimulatory ligands, T cell-activating cytokines and T cell-recruiting chemokines in these neutrophils, further reinforcing their antigen-presenting potential **(Fig. 3h)**. In contrast, MHC class II subunit expression remained unchanged **(Fig. S12F**).

To verify MHC class I antigen presentation by Nano ZSA-51D-reprogrammed neutrophils, we performed flow cytometry analysis on bone marrow neutrophils treated with Nano ZSA-51D and OT-I peptide (CSSSIINFEKL from OVA). Nano ZSA-51D significantly upregulated MHC class I expression on neutrophils (**Fig. 3i**), while MHC class II expression remained unchanged (**Fig. 3j**). Importantly, MHC class I presentation of OT-I peptide (H-2Kᵇ OT-I complex) was greatly enhanced 3.6-fold on neutrophils after Nano ZSA-51D treatment (**Fig. 3k**). Both immature/activated (CD101⁻CD14⁺) and mature/activated (CD101⁺CD14⁺) neutrophils demonstrated superior antigen presenting capacity compared to vehicle-treated control (**Fig. 3l**). These results suggest that Nano ZSA-51D reprograms neutrophils with enhanced MHC I antigen-presentation capacity, potentially enabling effective priming of CD8⁺ T cell immunity (**Fig. 3m**).

### Nano ZSA-51D-Reprogrammed Neutrophils Primes Antigen-Specific CD8^+^ T Cell Responses

To investigate whether Nano ZSA-51D-reprogrammed neutrophils can effectively prime antigen-specific CD8^+^ T cell activation and proliferation via MHC I-mediated antigen presentation, we conducted a co-culture experiment using OT-I CD8⁺ T cells from transgenic OT-I mice. We sorted total neutrophils from bone marrow cells that pre-treated with ovalbumin (OVA, 0.2 mg/mL) and Nano ZSA-51D (100 nM), and co-incubated them with OT-I CD8⁺ T cells for 48 hours (**Fig. 4a**).

**Figure 4.**
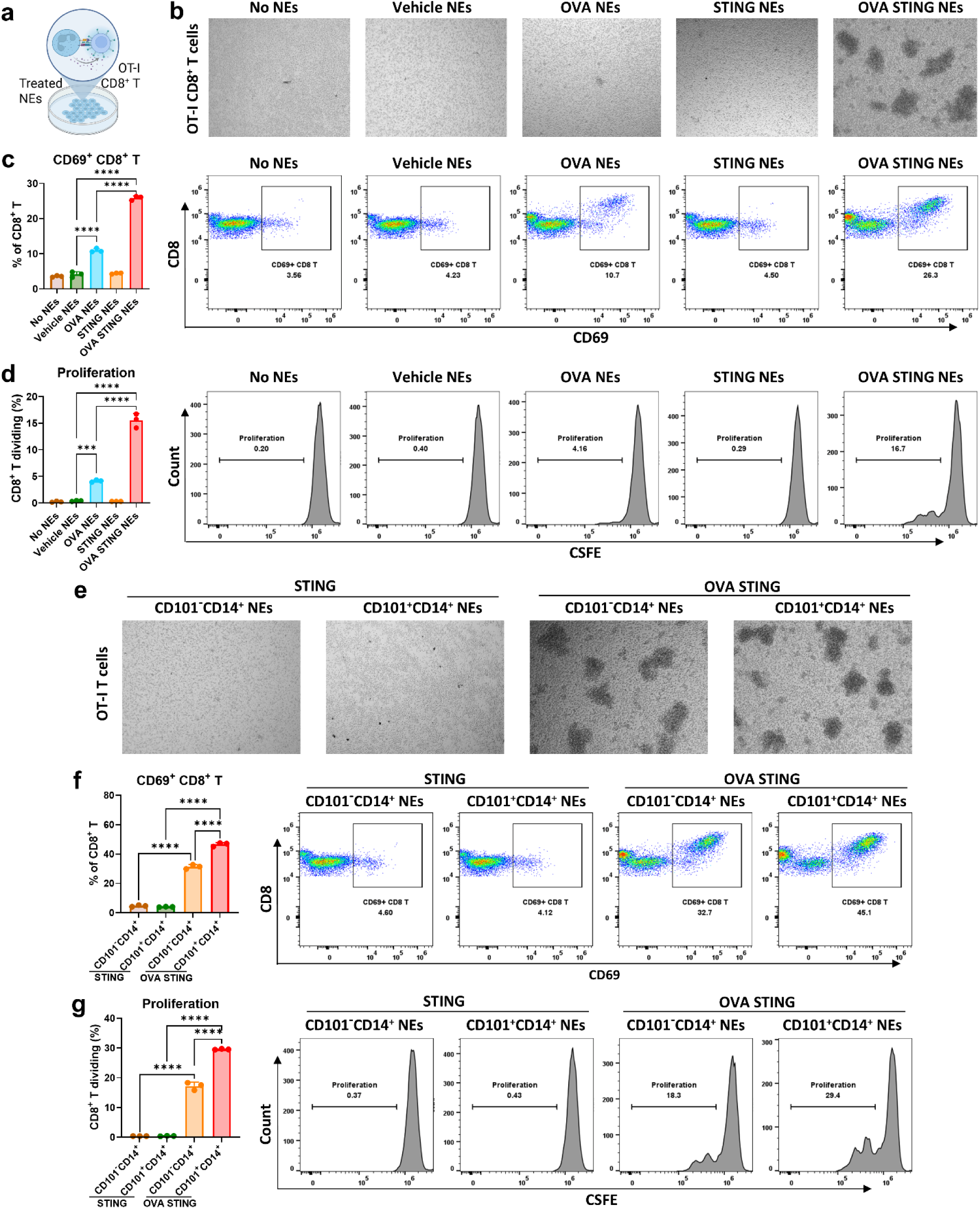
Nano ZSA-51D-reprogrammed neutrophils prime antigen-specific CD8^+^ T cell responses. **(a)** Experimental schematic showing Nano ZSA-51D (100 Nm) and OVA (0.2 mg/ml) pre-treated neutrophils (NEs) co-incubated with OT-I CD8⁺ T cells for 48 hours. **(b)** Bright-field microscopy images of OT-I CD8^+^ T cells after co-incubation with differently treated bone marrow neutrophils (NEs) for 48 hours. Neutrophils were sorted from bone marrow pretreated overnight with OVA (0.2 mg/ml) or/and Nano ZSA-51D (100 nM). **(c)** Quantification (left) and representative flow cytometry plots (right) of CD69^+^ OT-I CD8^+^ T cell activation after co-incubation with differently treated neutrophils for 48 hours (n = 3). ****p < 0.0001. **(d)** Quantification (left) and representative flow cytometry plots (right) of OT-I CD8^+^ T cell proliferation by CSFE staining after co-incubation with differently treated neutrophils for 48 hours (n=3). ***p < 0.001, ****p < 0.0001. **(e)** Bright-field microscopy images of OT-I CD8+ T cells after co-incubation with differently treated IA (CD101^-^CD14^+^) and MA (CD101^+^CD14^+^) neutrophils for 48 hours. IA (CD101^-^CD14^+^) and MA (CD101^+^CD14^+^) neutrophils were sorted from bone marrow pretreated overnight with Nano ZSA-51D (100 nM) with or without OVA. **(f)** Quantification (left) and representative flow cytometry plots (right) of CD69^+^ OT-I CD8^+^ T cell activation after co-incubation with differently treated IA (CD101-CD14+) and MA (CD101+CD14+) neutrophils for 48 hours (n=3). ****p < 0.0001. **(g)** Quantification (left) and representative flow cytometry plots (right) of OT-I CD8^+^ T cell proliferation by CSFE staining after co-incubation with differently treated IA (CD101-CD14+) and MA (CD101+CD14+) neutrophils for 48 hours (n=3). ****p < 0.0001.

Strikingly, neutrophils treated with OVA and Nano ZSA-51D induced extensive OT-I CD8^+^ T cell clustering (**Fig. 4b**), suggesting enhanced recruitment, activation, and proliferation of OT-I CD8^+^ T cells. Flow cytometry analysis further confirmed that OVA and Nano ZSA-51D co-treated neutrophils strongly activated OT-I CD8^+^ T cells, as indicated by a significant increase in CD69, CD25 and CD137 (4-1BB) expression (**Fig. 4c and Fig. S13A, B**). Moreover, they significantly promoted the proliferation of OT-I CD8⁺ T cells, as demonstrated by flow cytometry proliferation assays (**Fig. 4d**). These data suggest Nano ZSA-51D-reprogrammed neutrophils activate CD8^+^ T cell responses through enhanced MHC I antigen presentation.

To further assess the CD8⁺ T cell activation potential of immature and mature CD14⁺ neutrophils, we sorted these two distinct neutrophil subtypes from bone marrow cells that were treated with OVA and Nano ZSA-51D, and co-incubated them with OT-I CD8⁺ T cells for 48 hours. Both immature/activated (CD101⁻CD14⁺) and mature/activated (CD101⁺CD14⁺) neutrophils treated with OVA and Nano ZSA-51D induced prominent OT-I CD8^+^ T cell clustering (**Fig. 4e**). Further Flow cytometry analysis revealed that both CD101^-^CD14^+^ and CD101⁺CD14⁺ neutrophils strongly activated OT-I CD8⁺ T cells, as indicated by increased expression of CD69, CD25, and CD137 (4-1BB) (**Fig. 4f and Fig. S13C, D**) and significantly promoted OT-I CD8⁺ T cell proliferation (**Fig. 4g**). These results demonstrated that Nano ZSA-51D-reprogrammed immature and mature CD14^+^ neutrophils mediated potent antigen-specific CD8⁺ T cell activation and proliferation.

### Nano ZSA-51D in Combination with α-PD1 Achieved Potent Therapeutic Efficacy in Colon and Pancreatic Cancer Models

Next, we evaluated the antitumor efficacy of Nano ZSA-51D in both MC-38 colon cancer and pancreatic cancer model (**Fig. 5a**). Notably, Nano ZSA-51D (1 mg/kg, I.V.) in combination with α-PD1 (100 μg/mice) achieved 100% tumor complete remission (100% CR) without recurrence, leading to 100% long-term survival (**Fig. 5b, c, Fig. S14**). By comparison, free ZSA-51D with α-PD1 significantly inhibited tumor growth with 80% CR, while clinically tested STING agonist diABZI with 60% CR.

**Figure 5.**
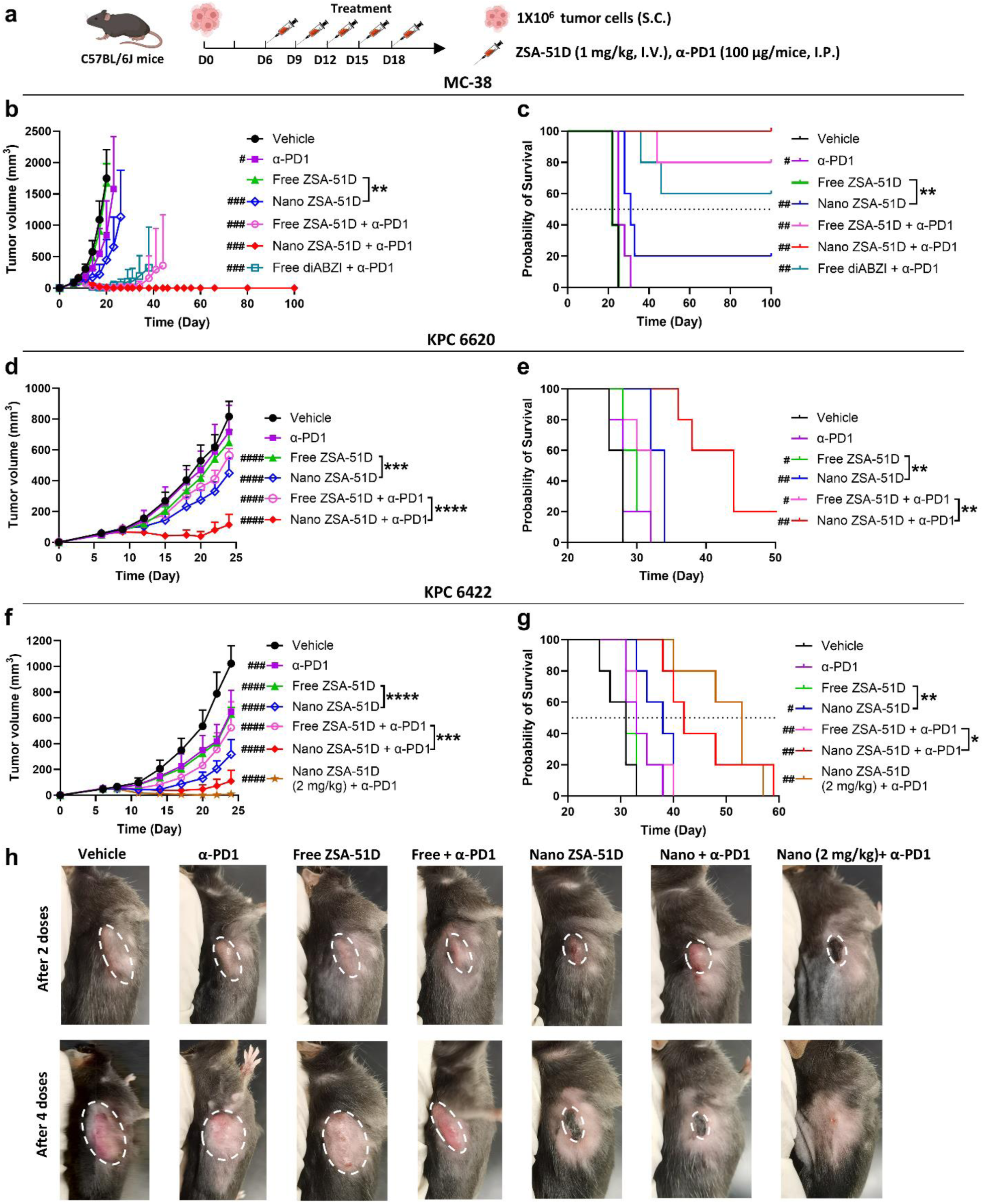
Nano ZSA-51D in combination with α-PD1 achieves potent therapeutic efficacy in colon and pancreatic cancer models. **(a)** Schematic Illustration of the cancer model establishment in C57BL/6J mice and treatment regimen in tumor-bearing mice. **(b-g)** Tumor growth curves (b, d, f) and survival curves (c, e, g) of MC-38 colon (b, c), KPC 6620 (d, e) and KPC 6422 (f, g) pancreatic cancer model following the indicated treatment regimens. Unless otherwise indicated, ZSA-51D or diABZI was administered at 1 mg/kg (I.V.) and α-PD1 at 100 μg/mouse (I.P.) (n = 5). *p < 0.05, **p < 0.01, ***p < 0.001, ****p < 0.0001. #p < 0.05, ##p < 0.01, ###p < 0.001, ####p < 0.0001 vs. vehicle group. **(h)** Representative tumor images of the KPC 6422 pancreatic cancer model at 2 and 4 doses post-treatment. The white dashed line indicates tumor margin.

Furthermore, we evaluated the antitumor efficacy of Nano ZSA-51D in pancreatic ductal adenocarcinoma (PDAC) which is not response to PD-1 therapy. We used KPC 6422 and KPC 6620 cell lines, derived from the KPC (Kras^G12D, p53^R172H, Pdx1-Cre) genetically engineered mouse model of PDAC.^55^ In both KPC 6422 and KPC 6620 pancreatic cancer models, Nano ZSA-51D (1 mg/kg, I.V.) with/without α-PD1 significantly reduced tumor burden and extend median survival, demonstrating superior efficacy over free ZSA-51D with/without α-PD1 (**Fig. 5d-g and Fig. S15, 16**). The Nano ZSA-51D at 2 mg/kg dose with α-PD1 led to significant tumor remission and extended median survival by 3 weeks compared to PD-1 monotherapy (**Fig. 5f and g**). In the KPC 6422 model, significant tumor shrinkage and scabbing were observed after the 4^th^ dose of Nano ZSA-51D (1 mg/kg, I.V.) with/without α-PD1 (**Fig. 5h**). In the 2 mg/kg Nano ZSA-51D plus α-PD1 group, tumor scabbing occurred after the 2^nd^ doses and completely disappeared after the 4^th^ dose, demonstrating rapid and effective tumor elimination.

### Nano ZSA-51D-Induced Neutrophil Infiltration Enhances Durable CD8^+^ T Cell Responses and Long-Term Memory in TME

To further examine whether Nano ZSA-51D with/without α-PD1 therapy induces neutrophil infiltration that enhances the CD8^+^ T cell-mediated antitumor immune responses in TME, we analyzed tumor-infiltrating neutrophils and T cells in MC-38 tumor-bearing mice by flow cytometry at 1- and 4-day post-treatment (**Fig. 6a**).

**Figure 6.**
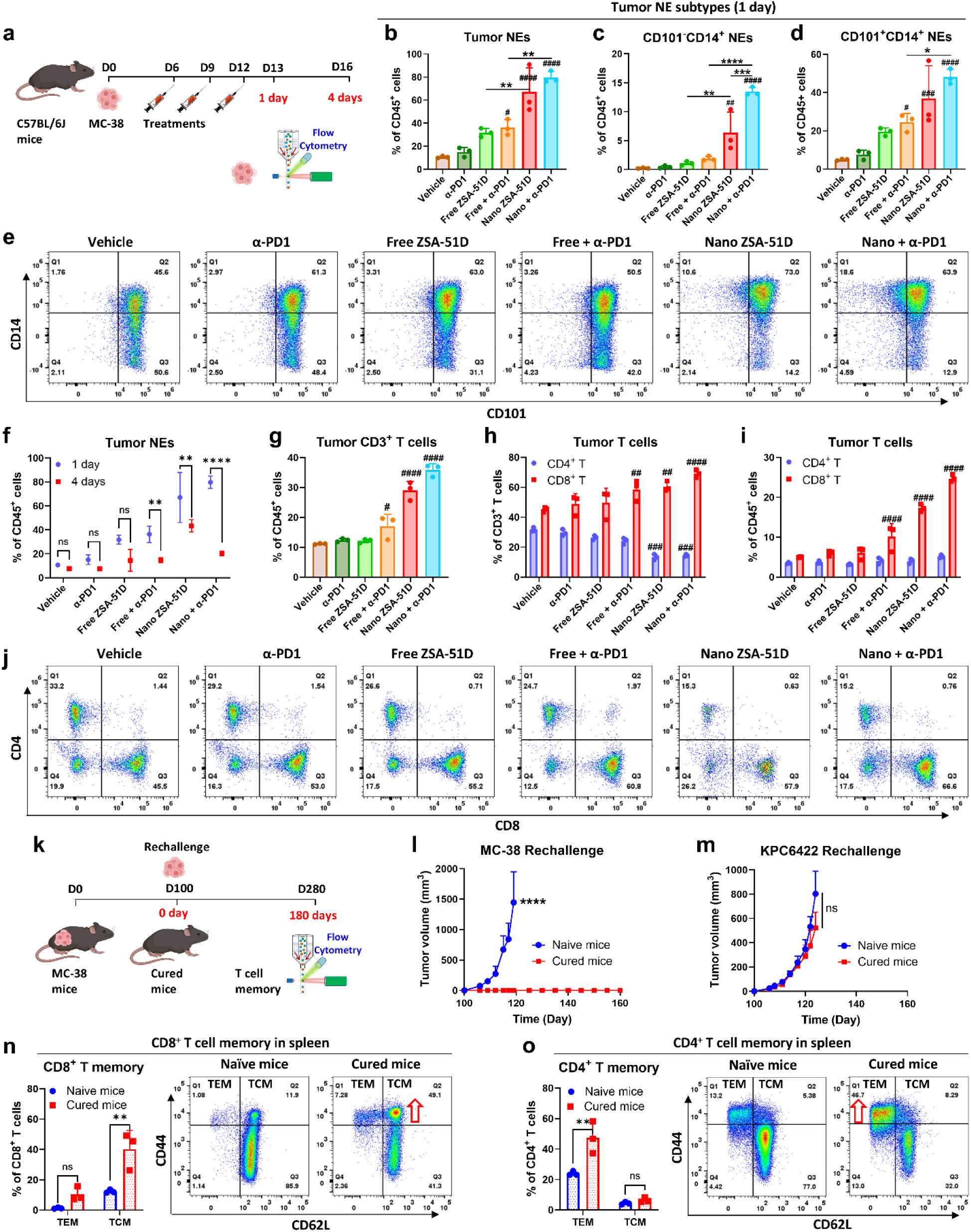
Nano ZSA-51D-induced neutrophil infiltration enhances durable CD8^+^ T cell responses and long-term memory in TME. **(a)** Schematic illustrating the treatment schedule and flow cytometry analysis of tumor-infiltrating immune cells in MC-38 tumor-bearing C57BL/6J mice at 1 day and 4 days post-treatment. **(b-d)** Quantification of total tumor-infiltrating neutrophils (NEs) (b), CD101^-^CD14^+^ (c) and CD101^+^CD14^+^ (d) NEs within CD45^+^ immune cells at 1-day post-treatment (n = 3). *p < 0.05, **p < 0.01, ***p < 0.001, ****p < 0.0001; #p < 0.05, ##p < 0.01, ###p < 0.001, ####p < 0.0001 vs. vehicle group. **(e)** Representative flow cytometry plots of CD101^-^CD14^+^ and CD101^+^CD14^+^ neutrophils in MC-38 tumors at 1-day post-treatment. **(f)** Quantification and comparison of tumor-infiltrating neutrophils within CD45^+^ immune cells at 1 day and 4 days post-treatment (n = 3). **p < 0.01, ****p < 0.0001. **(g-i)** Quantification of CD3^+^ T cells within CD45^+^ immune cells (g), CD4^+^ and CD8^+^ T cells within CD3^+^ T cells (h) or within CD45^+^ immune cells (i) at 4 days post-treatment (n = 3). #p < 0.05, ##p < 0.01, ###p < 0.001, ####p < 0.0001 vs. vehicle group. **(j)** Representative flow cytometry plots of tumor-infiltrating CD4^+^ and CD8^+^ T cells within CD3^+^ T cells at 4 days post-treatment. **(k)** Schematic of MC-38 tumor cell rechallenge in mice cured with Nano ZSA-51D and α-PD1 combination therapy at 100 days post-treatment. Long-term T cell memory was evaluated by flow cytometry at 180 day post-rechallenge. (**l, m**) Tumor growth curves following rechallenge with MC-38 (l) or KPC 6422 (m) tumor cell in naïve C57BL/6J mice and cured mice at 100 days post-treatment (n=5). ****p < 0.0001; ns, not significant. **(n, o)** Quantification (left) and representative flow cytometry plots (right) of CD8^+^ (n) and CD4^+^ (o) effector memory (TEM: CD62L^-^CD44^+^) and central memory (TCM: CD62L^+^CD44^+^) T cells in the spleen of naïve C57BL/6J mice and cured mice at 180-day post-rechallenge (n=3). **p < 0.01; ns, not significant.

At 1-day post-treatment, Nano ZSA-51D with α-PD1 dramatically increased tumor-infiltrating neutrophils from 10.6% to 79.7% of CD45^+^ immune cells, compared to vehicle-treated groups (**Fig. 6b and Fig. S17A**), as well as the immature/activated (CD101^-^CD14^+^) neutrophils from 0.22% to 13.5% (**Fig. 6c, e**), and the mature/activated (CD101^+^CD14^+^) neutrophils from 5% to 48.2% (**Fig. 6d, e**). In contrast, free ZSA-51D with/without α-PD1 also slightly enhanced total neutrophils and mature/activated (CD101^+^CD14^+^) neutrophils in tumors compared to vehicle-treated groups but did not significantly increase immature/activated (CD101^-^CD14^+^) neutrophils (**Fig. 6c-e)**. Conversely, 1-day post-treatment of Nano ZSA-51D with α-PD1 decreased macrophages from 59.7 % to 9.1 % of CD45^+^ immune cells (**Fig. S17B, C**). A similar trend was observed in the KPC 6620 pancreatic cancer model, where 1-day post-treatment of Nano ZSA-51D with α-PD1 increased tumor-infiltrating neutrophils from 31.9% to 63.3% (**Fig. S18A, C**) and decreased macrophages from 55.1% to 21.7% (**Fig. S18B, D**).

However, at 4-day post-treatment, tumor-infiltrating neutrophils declined compared to 1-day post-treatment, with the most substantial reduction observed from 79.7% to 20.4% in the Nano ZSA-51D with α-PD1 (**Fig. 6f and Fig. S19A**).

Concurrently, at 4-day post-treatment, Nano ZSA-51D with/without α-PD1 significantly increased tumor-infiltrating CD3^+^ T cells, with the most dramatic increase from 11.1% to 35.9% of CD45^+^ immune cells in Nano ZSA-51D with α-PD1 group, which is significantly higher than that of free ZSA-51D with/without α-PD1 (**Fig. 6g and Fig. S19B**). Further analysis revealed that Nano ZSA-51D with/without α-PD1 expanded tumor-infiltrating CD8⁺ T cells, while tumor-infiltrating CD4⁺ T cell proportions remained unchanged among CD45⁺ immune cells **(Fig. 6h-j)**. These results suggested that Nano ZSA-51D-induced infiltration of immature and mature activated CD14^+^ neutrophils enhances CD8⁺ T cell responses in the TME.

To assess whether Nano ZSA-51D with α-PD1 therapy generates tumor-specific immune memory, we performed tumor rechallenge experiments in cured MC-38 mice 100 days post-treatment (**Fig. 6k**). MC-38 tumor cells failed to establish tumors in Nano ZSA-51D with α-PD1 cured MC-38 mice, while KPC 6422 tumors grew at similar rates in both naïve and cured MC-38 mice (**Fig. 6l, m**). These results suggest that Nano ZSA-51D with α-PD1 therapy induces a tumor-specific immune memory response against MC-38 tumor cells but not against KPC 6422 tumor cells.

To further characterize the immune memory response, we also analyzed CD8⁺ and CD4⁺ T cell memory subsets in the spleen, lymph nodes and blood of cured MC-38 mice 180 days after tumor rechallenge (**Fig. 6k**). Flow cytometry analysis revealed a significant increase in CD8⁺ T central memory (TCM) cells in cured MC-38 mice, with a 3.3-, 4.4- and 4.5-fold increase in spleen, lymph nodes and blood compared to naïve mice (**Fig. 6n and Fig. S20A, C**). While CD4^+^ TCM cells had no significant change (**Fig. 6o and Fig. S20B, D**). These results further confirm that Nano ZSA-51D with α-PD1 therapy also generates the long term CD8^+^ T cell memory, which contributes to complete tumor remission and long-term tumor immunity.

## DISCUSSION

Although the immunosuppressive role of MDSCs in cancer has been extensively studied over the past decade, developing strategies to overcome neutrophil-mediated immune suppression remains challenging.^1,2,7^ Neutrophils exhibit remarkable plasticity and context-dependent functions, rapidly adapting to tumor-derived signals, which complicates efforts to define their roles and develop effective interventions against MDSC-mediated immune suppression.^3–6^

Recent evidence suggests that the immunosuppressive functions of MDSCs may be programmed as early as the hematopoietic stem and progenitor cell (HSPC) stage by suppressing interferon signaling.^20–25^ Given that downregulated interferon signaling underlies MDSC-mediated immunosuppression, we found that STING activation can activate STING-interferon signaling in HSPCs and reprogram them toward antitumor neutrophils to enhance cancer immunotherapy. In this study, we developed an albumin-STING nanoagonist (Nano ZSA-51D) that expanded HSPCs and reprogrammed them to generate antitumor neutrophils with enhanced MHC I antigen presentation, thereby activating CD8^+^ T cell-mediated antitumor immunity. We further demonstrated that Nano ZSA-51D combined with α-PD1 achieved potent anticancer efficacy in colon and pancreatic cancer models.

While prior studies have mainly focused on STING-induced activation of dendritic cells and macrophages, our work significantly broadens this paradigm by reprograming HSPCs to antitumor neutrophils as key effector cells in STING-based immunotherapy. Our findings reveal that Nano ZSA-51D expands HSPCs and skews their differentiation toward granulocyte-monocyte progenitors (GMP) through STING–NF-κB–IL-6 signaling, eventually expanding the neutrophil lineage. This suggests that STING signaling in the bone marrow is not merely an immune-sensing pathway but also a potent regulator of hematopoietic fate of HSPCs. Mechanistically, STING activation driven NF-κB-IL-6 signaling favors granulocyte-monocyte lineage commitment. Indeed, IL-6 has been reported to drive HSPC proliferation and differentiation toward granulocyte-monocyte progenitors that promote granulopoiesis, aligning with our observation.^56–58^ In addition, Nano ZSA-51D activates STING-interferon signaling in HSPCs and reprograms their transcriptional landscape toward an antitumor immune phenotype, thereby promoting differentiation into antitumor neutrophils rather than MDSCs. This finding suggests that early STING activation at the progenitor level can reshape myelopoiesis, restoring immune surveillance and preventing the pathological programming of immunosuppressive myeloid lineages by cancer-derived signals.

Furthermore, this study reveals a previously underexplored antitumor role of neutrophils in STING-mediated cancer immunotherapy.^59–61^ We demonstrate that both immature (CD101⁻) and mature (CD101⁺) bone marrow neutrophils are reprogrammed into CD14⁺ICAM-1^+^ subsets via STING–NF-κB–TNF-α signaling by Nano ZSA-51D in bone marrow. These CD14⁺ICAM-1^+^ neutrophils infiltrate tumors more effectively and exhibit potent synergy with anti-PD1 therapy. Notably, adoptive transfer of Nano ZSA-51D-reprogrmamed immature CD14⁺ neutrophils in combination with α-PD1 leads to 100% complete tumor remission through long-term CD8⁺ T cell central memory, highlighting their potent anti-tumor immune function. The upregulation of CD14 and ICAM-1 on neutrophils likely underlies these antitumor functions. CD14 acts as a co-receptor for toll-like receptors, activating innate immunity responses to pathogens by binding PAMPs and DAMPs.^44,62–64^ While neutrophil ICAM-1 (CD54) expression is correlated with enhanced migration, phagocytosis and costimulatory signal of T cell activation.^46,47,65–67^ Studies also suggest that ICAM-1 preferentially co-stimulates CD8^+^ T cells while providing weak co-stimulation to CD4^+^ T cells.^68,69^ Together, activation of CD14 and ICAM-1 on neutrophils may enhance their responsiveness to PAMPs and DAMPs within the tumor microenvironment, promoting tumor infiltration, antigen presentation, and activation of innate and adaptive immunity in synergy with immune checkpoint blockade.

Transcriptomic and immune profiling reveal that Nano ZSA-51D-reprogrammed CD14⁺ neutrophils upregulate interferon signaling and display high MHC class I levels, along with enhanced costimulatory ligands and signaling. This confers them with enhanced MHC class I antigen-presentation capacity, thereby supporting antigen-specific CD8^+^ T cell-mediated antitumor immunity. Previous studies also identified a subset of ICAM-1^hi^CD14^+^MHC I^hi^ APC-like “hybrid neutrophils”, derived from immature progenitors, that are capable of cross-presenting antigens and initiating robust anti-tumor T cell responses in early-stage human lung cancer.^38,39,70^ In contrast, CD11b^+^CD14^-^ MDSCs have been shown to exert immunosuppressive effects on CD8⁺ T cells, contributing to immune evasion in non-small cell lung cancer.^40^ Collectively, these findings suggest that Nano ZSA-51D induces CD14^+^ICAM-1^+^MHC I^hi^ neutrophils, acquiring antitumor properties through enhanced MHC I-mediated antigen presentation to prime CD8^+^ T cell responses.

Finally, despite extensive efforts to develop STING agonists that mimic natural cyclic dinucleotide (CDNs), their clinical efficacy has remained limited.^71–73^ CDN-based STING agonists suffer from rapid enzymatic degradation and poor cellular permeability due to their hydrophilic and anionic nature, restricting their use to intratumoral administration. To overcome these limitations, nanoparticle-formulated CDNs and non-CDN STING agonists with improved pharmacological properties have been developed for systemic bioavailability, demonstrating potent antitumor activity.^33,73–77^ Nevertheless, these non-CDN STING agonists may still have limited capacity to reprogram HSPCs within the bone marrow and are often associated with toxicities. In contrast, our albumin-STING nanoagonist, Nano ZSA-51D, exhibits unique anticancer efficacy by reprogramming HSPCs into antitumor neutrophils in the bone marrow, while maintaining a favorable safety profile. We evaluated the toxicity profile of Nano ZSA-51D by using single and multiple doses (5 doses, 1 mg/kg, I.V.). Complete blood cell count, liver functions, and histopathological assessments revealed no significant toxicity (**Fig. S21**), indicating that Nano ZSA-51D is well tolerated.

In summary, we demonstrate that albumin-STING nanoagonist, Nano ZSA-51D, expands HSPCs and reprograms them into antitumor neutrophils in the bone marrow, which subsequently migrate into TME to enhance MHC I-mediated antigen presentation and activate CD8^+^ T cell antitumor immunity. Nano-ZSA51D combined with α-PD1 achieves potent anticancer efficacy in both colon and pancreatic cancer models. These data suggest that Nano ZSA-51D provides a novel strategy to reprogram HSPCs into antitumor neutrophils and highlight the potential of early interventions at HSPC stage to rewire neutrophil fate for effective immunotherapy.

## MATERIALS AND METHODS

### Cell cultures

All cells were maintained at 37 °C in a humidified incubator (5% CO₂, 95% air). THP-1-Blue™ ISG cells (InvivoGen) were cultured in RPMI 1640 with 10% heat-inactivated FBS and 100 µg/ml normocin. LSK cells from WT C57BL/6J mice were maintained in StemSpan medium with 50 ng/ml mouse SCF (BioLegend). Bone marrow neutrophils from WT and STING KO mice (C57BL/6J-Sting1gt/J) and CD8⁺ T cells from OT-I mice were cultured in RPMI 1640 with 10% FBS. MC-38 colon adenocarcinoma cells (Kerafast) were cultured in DMEM with 10% FBS and gentamycin. KPCY pancreatic cancer cells (KPC 6620 and KPC 6422; Kerafast) were cultured in DMEM with GlutaMAX, 10% FBS, and sodium pyruvate. Unless otherwise indicated, all media contained standard supplements (L-glutamine, HEPES, sodium pyruvate, nonessential amino acids, and Pen/Strep). Key reagents and resources are listed in **Table S1**.

### Mice and tumor models

All animal experiments were approved by the University of Michigan UCUCA and performed under pathogen-free, temperature- and humidity-controlled conditions with a 12 h light/dark cycle. WT C57BL/6J mice (Charles River), STING KO mice (C57BL/6J-Sting1gt/J), and OT-I transgenic mice (C57BL/6-Tg(TcraTcrb)1100Mjb/J) (Jackson Laboratory) aged 6–8 weeks were used. For tumor studies, 1×10⁶ MC38, KPC 6620, or KPC 6422 cells in 100 µL FBS-free DMEM were injected subcutaneously into the right flank of 6-week-old male WT mice. Mice were randomized for treatment when tumors reached 50–100 mm³ (day 6 post-inoculation), and tumor growth and survival were monitored every 2–3 days. Tumor volume was calculated as length × width²/2.

### Synthesis and characterization of dimeric STING agonist ZSA-51D

The dimeric STING agonist ZSA-51D were synthesized based on our previously reported monomeric STING agonist ZSA-51 (**Fig. S1**).^33,76^ Briefly, starting material **1** was suspended in Et_2_O and cooled to 0°C. MeMgBr was added to the suspension dropwise. The resulting mixture was stirred overnight to give intermediate **2**. **2** was stirred with PCC in DCM overnight to afford intermediate **3**. The solution of **3**, DIPEA and methyl thioglycolate in DMA was stirred at room temperature for 40 min under nitrogen atmosphere, followed by the addition of *t*-BuOK. The resulting mixture was heated at 80°C for 14 h to afford intermediate **4**. **4** was stirred with HATU, DCM, DIPEA and methyl 3-aminopropionate hydrochloride or *tert*-butyl 3-aminopropionate hydrochloride for 2 h at room temperature to provide intermediate **5a** or **5b**. **5a** or **5b** was deprotected under hydrogen atmosphere with palladium catalyst in ethanol to give intermediate **6a** or **6b**. **6a** or **6b** was dissolved in DCM. Then, TBDMSCl, imidazole and DMAP were added. The solution was stirred at room temperature for 0.5 h to give intermediate **7a** or 7**b**. The mixture of **7a** or **7b**, NBS, AIBN and CCl_4_ was heated at 80°C for 2 h to produce intermediate **8a** or **8b**. **8a** or **8b** was suspended in CH_3_CN and cooled to 0°C. Then, NMO was added. The mixture was stirred at room temperature overnight to give intermediate **9a** or **9b**. **9a** or **9b** was dissolved in DCM followed by the addition of PCC. The suspension was stirred at room temperature for 16 h to furnish intermediate **10a** or **10b**. TBAF (0.25 mL, 1 M in THF) was added into the solution of **10a** or **10b** in THF. The solution was stirred at room temperature for 0.5 h to afford intermediate **11a** or **11b**. The mixture of **11a** or **11b**, K_2_CO_3_, catalytic amount of NaI, CH_3_CN and 1,3-dibromopropane was heated at 80°C for 3 h to produce intermediate **12a** or **12b**. **12a** or **12b** was mixed **11a** or **11b**, K_2_CO_3_, catalytic amount of NaI and CH_3_CN, and heated at 80°C for 4 h to provide target compound ZSA-51D or intermediate **13b**. **13b** was dissolved in DCM followed by the addition of TFA. The solution was stirred at room temperature for 3 h to afford target compound ZSA-52D.

### hSTING protein binding assay

The human STING WT binding assay (Revvity, PerkinElmer) was performed in white 386-well plates per the manufacturer’s instructions. Test compounds (5 µL) were mixed with human STING WT protein (5 µL, 6×His-tagged), followed by 10 µL of premixed d2 ligand and Tb cryptate-conjugated antibody. Reactions were incubated at room temperature for 3 h, and HTRF signals were measured using a Synergy 2 microplate reader (BioTek).

### THP1-Blue ISG cell assay

THP1-Blue ISG cells (InvivoGen), a human monocytic line expressing a SEAP reporter under an interferon-stimulated response element, were used to monitor IRF activation. Cells (1×10⁶/mL) were incubated with test compounds (100 µL) for 24 h at 37 °C. Supernatants (20 µL) were mixed with 180 µL QUANTI-Blue™ reagent (InvivoGen) in 96-well plates and incubated for 2 h at 37 °C. SEAP activity was measured at 620 nm using a Synergy 2 microplate reader (Biotek). All assays were run in triplicate with vehicle controls.

### Nano ZSA-51D preparation

Nano ZSA-51D was prepared as previously described.^35^ Briefly, 10 mg ZSA-51D in 1 mL chloroform was emulsified with 100 mg mouse serum albumin in 20 mL Milli-Q water using a rotor-stator homogenizer. The emulsion was processed through six cycles at 20,000 psi with a high-pressure homogenizer (4 °C), and chloroform was removed under reduced pressure at 25 °C. The resulting suspension was lyophilized and stored at −20 °C. Particle size was measured by dynamic light scattering (DLS; Zetasizer Nano-ZS, Malvern), with three measurements per sample and experiments performed in triplicate.

### Western blot analysis of STING pathway activation

Bone marrow cells from C57BL/6 mice were treated with vehicle, free, or Nano ZSA-51D (10 or 100 nM) for 4 h, lysed in RIPA buffer with protease and phosphatase inhibitors, sonicated, and centrifuged at 12,000 × g for 15 min at 4 °C. Protein concentrations were determined by BCA assay. Equal amounts of protein were separated by SDS-PAGE, transferred to PVDF membranes, blocked with 5% milk in TBST, and incubated with primary antibodies against STING, p-STING (Ser365), IRF3, p-IRF3 (Ser396), NF-κB p65, and p-NF-κB p65 (Ser536), followed by HRP-conjugated secondary antibodies. Signals were detected using ECL (Bio-Rad) and imaged with a ChemiDoc MP system.

### Cytokine secretion assay

Bone marrow cells were treated with vehicle, free ZSA-51D, or Nano ZSA-51D (100 nM) for 48 h. Supernatants were collected, and IFN-β, TNF-α, and IL-6 levels were quantified using mouse-specific ELISA kits (R&D Systems) per the manufacturer’s instructions. Absorbance was measured at 450 nm (Synergy 2, Biotek), and cytokine concentrations were calculated from standard curves.

### In vivo treatment regimen

Mice were intravenously administered free or Nano ZSA-51D (1 mg/kg), free diABZI (1 mg/kg), or vehicle (15 mg/kg mouse serum albumin). Free ZSA-51D or diABZI was dissolved in DMSO/PEG-400 (1:4, v/v) and diluted 1:1 with PBS to 0.2 mg/mL for injection; Nano ZSA-51D was directly resuspended in PBS at 0.2 mg/mL. For combination therapy, 100 μg α-PD1 antibodies (Bio X Cell) were given intraperitoneally according to the experimental schedule. For antitumor efficacy, treatments were given every 3 days for 5 doses; for HSPC and immune profiling, treatments were given every 3 days for 3 doses.

### In vivo PK study

Free or Nano ZSA-51D (1 mg/kg) was administered intravenously to MC-38 tumor-bearing C57BL/6J mice. Blood and tissues were collected at 15 min, 2 h, and 7 h post-injection. Plasma was separated by centrifugation (15,000 rpm, 10 min). After euthanasia, tumors and hind limb bones were harvested; bone marrow was isolated by brief centrifugation. Samples were stored at −80 °C until analysis. ZSA-51D and ZSA-52D levels were quantified by LC–MS/MS using a Waters XBridge C18 column and a Triple Quad 5500 mass spectrometer (SCIEX) with ESI in positive mode, employing analyte-specific MRM transitions. Results are reported as ng/mL (plasma) or ng/g (tissue).

### Single-cell suspension preparation and flow cytometry analysis

Peripheral blood, bone marrow, and tumors were collected after STING agonist treatment. Blood (200 µL) was drawn from ophthalmic veins, and tumors and hind limb bones were harvested post-euthanasia. Bone marrow was flushed with ice-cold PBS + 1% FBS and 1 mM EDTA through a 70 µm strainer and centrifuged (350 × g, 5 min). Tumors (<1 g) were minced, digested in RPMI 1640 with 0.01–0.1% collagenase/hyaluronidase and 0.15 mg/mL DNase I at 37 °C for 30 min with shaking, filtered, and washed. All suspensions underwent RBC lysis (ACK buffer, 5 min, 4 °C), washed, centrifuged (400 × g, 5 min), and resuspended at 1×10⁷ cells/mL. Cells were stained with Ghost Dye™ Violet 510 (30 min, 4 °C), blocked with anti-CD16/32 (10 min, 4 °C), then stained with fluorochrome-conjugated antibodies (**Table S1**) for 20 min at 4 °C, washed, fixed (BioLegend), and analyzed on a Cytek Aurora cytometer; data were processed using FlowJo (**Fig. S22**).

### HSPC analysis

Bone marrow single-cell suspensions were enriched for lineage-negative (Lin⁻) cells using a biotin-conjugated lineage panel (BioLegend) and magnetic separation, then stained with fluorophore-conjugated antibodies (**Table S1**) for flow cytometric analysis of HSPC subsets (**Fig. S23**).

### Cell sorting by FACS

Neutrophils and subpopulations were isolated from bone marrow treated in vitro with vehicle or Nano ZSA-51D (100 nM) using a Sony MA900 sorter with a 100 μm chip at 4 °C. Cells were stained with CD11b, Ly6G, CD101, and CD14, and dead cells excluded with DAPI (5 µg/mL). Neutrophils (CD11b⁺Ly6G⁺) were further gated by CD101 and CD14 to yield CD101⁻CD14⁻, CD101⁻CD14⁺, and CD101⁺CD14⁺ populations. Sorted cells were used for in vitro assays, adoptive transfer, or bulk mRNA analysis. LSK cells (Lin⁻Sca-1⁺c-Kit⁺) were sorted from enriched bone marrow Lin⁻ cells.

### LSK culture and treatment

Sorted LSK cells were cultured in StemSpan with 50 ng/mL SCF (BioLegend) at 2×10⁴ cells/well in 96-well U-bottom plates. Cells were treated with 100 nM Nano ZSA-51D or 10 ng/mL cytokines (TNF-α, IFN-β, IFN-γ, IL-6) for 3 days, then collected, washed, and resuspended in PBS for flow cytometry or bulk mRNA-seq analysis.

### In vitro reprogramming of bone marrow neutrophils

Bone marrow cells were isolated from femurs and tibiae of WT or STING KO mice (6–8 weeks). Cells were flushed with PBS + 1% FBS and 1 mM EDTA, centrifuged, RBC-lysed, washed, and resuspended in complete RPMI 1640. Cells (5×10⁶/well) were seeded in 12-well plates and treated overnight with 100 nM free or Nano ZSA-51D, vehicle (30 µg/mL mouse serum albumin). In addition, the sorted bone marrow neutrophils were treated with 10 ng ml^−1^ cytokines (TNF-α, IFN-β, IFN-γ, IL-6, TGF-β) for overnight. For TNF-α neutralization, cells were pretreated with TNF-α antibody (10 μg/mL) 2 hours in advance of Nano ZSA-51D treatment. After treatment, cells were collected, washed, and resuspended in PBS for flow cytometry (**Fig. S24**).

### Adoptive transfer of neutrophil subsets

Sorted neutrophil subsets (CD101⁻CD14⁻, CD101⁻CD14⁺, CD101⁺CD14⁺; 2×10⁶ cells/mouse) from Nano ZSA-51D–treated bone marrow were adoptively transferred intratumorally into MC-38 tumors, and mice received intraperitoneal α-PD1 (100 μg) every 2 days. Tumor growth and survival were monitored every 2–3 days.

### Tumors rechallenge and T cell memory analysis

To assess long-term immunity, cured mice were rechallenged 100–120 days post-treatment with MC-38 or KPC 6642 cells (5×10^6^ cells/mL, 100 µL) subcutaneously, alongside age-matched naïve controls. Tumor growth was monitored every 2–3 days from day 6, and tumor volume was recorded. At 180 days post-rechallenge, spleens, lymph nodes, and blood were collected to prepare single-cell suspensions for flow cytometric analysis of CD4⁺ and CD8⁺ T cell memory (**Fig. S25**).

### Bulk mRNA-seq analysis

Total RNA was extracted (RNeasy Plus, QIAGEN), and samples with RIN ≥ 8 were used. Poly(A) mRNA (250 ng) was isolated (NEBNext Poly(A) Magnetic Isolation), converted to cDNA libraries (NEBNext UltraExpress RNA Library Prep with dual indices), pooled, and sequenced on Illumina NovaSeq X 10B (150 bp paired-end). Adapters and low-quality bases were trimmed with Cutadapt, quality checked with FastQC, and contamination screened with Fastq Screen. Reads were aligned to GRCm38 (ENSEMBL) using STAR, quantified with RSEM, and metrics summarized with MultiQC. Differential expression was analyzed using DESeq2; genes with adjusted p < 0.05 and |log₂FC| > 1 were considered significant. KEGG pathway ORA and GSEA using KEGG/GO gene sets were performed for functional enrichment.

### Antigen presentation by neutrophils

Bone marrow cells from WT C57BL/6J mice were seeded at 1×10⁶ cells/well in 24-well plates and treated overnight with OT-I peptide (SIINFEKL, 50 μg/mL), Nano ZSA-51D (100 nM), or vehicle (30 μg/mL mouse serum albumin) in complete RPMI 1640. Cells were collected, washed, and stained with antibodies. MHC I expression and OT-I peptide presentation (H-2Kb/SIINFEKL) on neutrophils and subsets were analyzed by flow cytometry.

### OT-I CD8+ T cell activation and proliferation

OT-I CD8⁺ T cells were isolated (EasySep CD8⁺ kit, STEMCELL), labeled with CellTrace™ Violet, and washed. Bone marrow neutrophils and subsets (CD101⁻CD14⁺, CD101⁺CD14⁺) pretreated with OVA (200 μg/mL), Nano ZSA-51D (100 nM), or vehicle were sorted and co-cultured with labeled T cells at a 2:1 neutrophil:T cell ratio for 48 h; T cells alone served as controls. Cells were imaged by bright-field microscopy (EVOS M5000) and analyzed by flow cytometry for activation markers (CD69, CD25, CD137) and proliferation via CellTrace dilution.

### In vivo toxicity assessment

Six-week-old WT C57BL/6J mice (n = 3 per group) received vehicle (15 mg/kg albumin), free, or Nano ZSA-51D (1 mg/kg) intravenously every 3 days for five doses. Blood was collected on days 0, 3, and 15 for complete blood counts, and plasma on day 15 for liver function tests. Major organs (heart, liver, spleen, lung, kidney, intestine) were harvested for histopathology. Analyses were performed by the Unit for Laboratory Animal Medicine (ULAM).

### Statistical analysis

All statistical analyses were performed using GraphPad Prism (version 10.3.1) unless otherwise specified. Data are presented as mean ± standard deviation (SD). Comparisons between two groups were performed using an unpaired two-tailed Student’s T-test. For comparisons involving more than two groups, one-way or two-way analysis of variance (ANOVA) followed by Tukey’s or Sidak’s multiple comparison test was used as appropriate. For survival analyses, Kaplan–Meier curves were plotted and compared using the log-rank (Mantel–Cox) test. A p-value < 0.05 was considered statistically significant. Specific statistical tests used for each experiment are detailed in the figure legends.

## List of Supplementary Materials

Fig. S1 to S25

Table S1

## Acknowledgements

We thank the University of Michigan Unit for Laboratory Animal Medicine (ULAM) for support in maintaining laboratory animals, and the In Vivo Animal Core (IVAC) for assistance toxicity studies. We also acknowledge the University of Michigan Pharmacokinetics Core, Flow Cytometry Core, Genomics Core for technical service.

## Author contributions

J.T., W.G., and D.S. conceived the project, developed the hypotheses, designed the experiments and wrote the manuscript; J.T. performed most experiments and analyzed data, with assistance from all authors; H.Z. conducted the chemical synthesis. C.L. provided support for bioinformatic analysis. H.W. and F.K. assisted with the in vivo flow cytometry studies. Q.L., M.H. and B.W. assisted with the in vivo pharmacokinetic studies. Z.L. tested the STING binding and cellular activity in vitro. D.S. supervised the project and acquired funding.

## Competing interests

University of Michigan has filed a patent based on these studies where some authors are listed as inventors.

## Data and materials availability

All data associated with this study are present in the paper or the Supplementary Materials. Materials can be made available through a material transfer agreement with D.S.

## Notes

### Competing Interest Statement

The authors have declared no competing interest.

### Summary of Updates

This version of the manuscript has been revised to clarify the objectives and context in the Abstract and Introduction. The order of the Figures has been reorganized to improve the logical flow and clarity of the study. In addition, new data have been added to further support the conclusions and strengthen the overall findings. Minor textual edits were made throughout the manuscript for improved readability and consistency.

## References

1 Shaul, M. E. & Fridlender, Z. G. Tumour-associated neutrophils in patients with cancer. Nat Rev Clin Oncol 16, 601–620 (2019). 10.1038/s41571-019-0222-4

2 Veglia, F., Sanseviero, E. & Gabrilovich, D. I. Myeloid-derived suppressor cells in the era of increasing myeloid cell diversity. Nat Rev Immunol 21, 485–498 (2021). 10.1038/s41577-020-00490-y

3 Hedrick, C. C. & Malanchi, I. Neutrophils in cancer: heterogeneous and multifaceted. Nat Rev Immunol 22, 173–187 (2022). 10.1038/s41577-021-00571-6

4 Jaillon, S., et al. Neutrophil diversity and plasticity in tumour progression and therapy. Nat Rev Cancer 20, 485–503 (2020). 10.1038/s41568-020-0281-y

5 Huang, X., et al. Neutrophils in Cancer immunotherapy: friends or foes? Mol Cancer 23, 107 (2024). 10.1186/s12943-024-02004-z

6 Zhang, M., Qin, H., Wu, Y. & Gao, Q. Complex role of neutrophils in the tumor microenvironment: an avenue for novel immunotherapies. Cancer Biol Med 21, 849–863 (2024). 10.20892/j.issn.2095-3941.2024.0192

7 Eruslanov, E., Nefedova, Y. & Gabrilovich, D. I. The heterogeneity of neutrophils in cancer and its implication for therapeutic targeting. Nat Immunol 26, 17–28 (2025). 10.1038/s41590-024-02029-y

8 Su, J. L., et al. Pretreatment neutrophil-to-lymphocyte ratio is associated with immunotherapy efficacy in patients with advanced cancer: a systematic review and meta-analysis. Sci Rep-Uk 15 (2025). https://doi.org:ARTN 44610.1038/s41598-024-84890-3

9 Mitchell, K. G., et al. Neutrophil expansion defines an immunoinhibitory peripheral and intratumoral inflammatory milieu in resected non-small cell lung cancer: a descriptive analysis of a prospectively immunoprofiled cohort. J Immunother Cancer 8 (2020). https://doi.org:ARTN e000405 10.1136/jitc-2019-000405

10 Gungabeesoon, J., et al. A neutrophil response linked to tumor control in immunotherapy. Cell 186, 1448–1464 e1420 (2023). 10.1016/j.cell.2023.02.032

11 Linde, I. L., et al. Neutrophil-activating therapy for the treatment of cancer. Cancer Cell 41, 356-+ (2023). 10.1016/j.ccell.2023.01.002

12 Li, M., Ng, M. & Ng, L. G. Firing up neutrophil anti-tumor immunity with cocktails. Cancer Cell 41, 227–229 (2023). 10.1016/j.ccell.2023.01.005

13 Hirschhorn, D., et al. T cell immunotherapies engage neutrophils to eliminate tumor antigen escape variants. Cell 186, 1432-+ (2023). 10.1016/j.cell.2023.03.007

14 Kumbhojkar, N., et al. Neutrophils bearing adhesive polymer micropatches as a drug-free cancer immunotherapy. Nat Biomed Eng 8 (2024). 10.1038/s41551-024-01180-z

15 Ager, A. Cancer immunotherapy: T cells and neutrophils working together to attack cancers. Cell 186, 1304–1306 (2023). 10.1016/j.cell.2023.03.005

16 Benguigui, M., et al. Interferon-stimulated neutrophils as a predictor of immunotherapy response. Cancer Cell 42 (2024). 10.1016/j.ccell.2023.12.005

17 Wu, Y., et al. Neutrophil profiling illuminates anti-tumor antigen-presenting potency. Cell 187, 1422–1439 e1424 (2024). 10.1016/j.cell.2024.02.005

18 Mysore, V., et al. FcγR engagement reprograms neutrophils into antigen cross-presenting cells that elicit acquired anti-tumor immunity. Nature Communications 12 (2021). https://doi.org:ARTN 479110.1038/s41467-021-24591-x

19 Ozel, I., et al. Neutrophil-specific targeting of STAT3 impairs tumor progression via the expansion of cytotoxic CD8(+) T cells. Signal Transduct Target Ther 10, 279 (2025). 10.1038/s41392-025-02363-z

20 Garner, H., et al. Understanding and reversing mammary tumor-driven reprogramming of myelopoiesis to reduce metastatic spread. Cancer Cell (2025). 10.1016/j.ccell.2025.04.007

21 Dong, L. & Ng, L. G. Tumor neutrophils rewired from birth. Cancer Cell (2025). 10.1016/j.ccell.2025.05.005

22 Giles, A. J., et al. Activation of Hematopoietic Stem/Progenitor Cells Promotes Immunosuppression Within the Pre-metastatic Niche. Cancer Research 76, 1335–1347 (2016). 10.1158/0008-5472.Can-15-0204

23 Daman, A. W., et al. Microbial cancer immunotherapy reprograms hematopoiesis to enhance myeloid-driven anti-tumor immunity. Cancer Cell (2025). 10.1016/j.ccell.2025.05.002

24 Qian, J., et al. A CXCR4 partial agonist improves immunotherapy by targeting immunosuppressive neutrophils and cancer-driven granulopoiesis. Cancer Cell 43, 1512–1529 e1511 (2025). 10.1016/j.ccell.2025.06.006

25 Hegde, S., et al. Myeloid progenitor dysregulation fuels immunosuppressive macrophages in tumours. Nature (2025). 10.1038/s41586-025-09493-y

26 Alicea-Torres, K., et al. Immune suppressive activity of myeloid-derived suppressor cells in cancer requires inactivation of the type I interferon pathway. Nat Commun 12, 1717 (2021). 10.1038/s41467-021-22033-2

27 Sun, Y. Y. et al. The downregulation of type I IFN signaling in G-MDSCs under tumor conditions promotes their development towards an immunosuppressive phenotype. Cell Death Dis 13 (2022). https://doi.org:ARTN 3610.1038/s41419-021-04487-w

28 Ishikawa, H. & Barber, G. N. STING is an endoplasmic reticulum adaptor that facilitates innate immune signalling. Nature 455, 674–U674 (2008). 10.1038/nature07317

29 Ishikawa, H., Ma, Z. & Barber, G. N. STING regulates intracellular DNA-mediated, type I interferon-dependent innate immunity. Nature 461, 788–U740 (2009). 10.1038/nature08476

30 Balka, K. R., et al. TBK1 and IKKε Act Redundantly to Mediate STING-Induced NF-κB Responses in Myeloid Cells. Cell Rep 31 (2020). https://doi.org:ARTN 10749210.1016/j.celrep.2020.03.056

31 Liao, W. N., Du, C. H. & Wang, J. P. The cGAS-STING Pathway in Hematopoiesis and Its Physiopathological Significance. Front Immunol 11 (2020). https://doi.org:ARTN 57391510.3389/fimmu.2020.573915

32 Xie, J. Y., et al. STING activation in TET2-mutated hematopoietic stem/progenitor cells contributes to the increased self-renewal and neoplastic transformation. Leukemia 37, 2457–2467 (2023). 10.1038/s41375-023-02055-z

33 Zhao, H. Y., et al. An oral tricyclic STING agonist suppresses tumor growth through remodeling of the immune microenvironment. Cell Chem Biol 32 (2025). 10.1016/j.chembiol.2025.01.004

34 Hoogenboezem, E. N. & Duvall, C. L. Harnessing albumin as a carrier for cancer therapies. Adv Drug Deliver Rev 130, 73–89 (2018). 10.1016/j.addr.2018.07.011

35 Song, Y. D., et al. Albumin nanoparticle containing a PI3Kγ inhibitor and paclitaxel in combination with α-PD1 induces tumor remission of breast cancer in mice. Sci Transl Med 14 (2022). https://doi.org:ARTN eabl364910.1126/scitranslmed.abl3649

36 Sun, D., Zhou, S. & Gao, W. What Went Wrong with Anticancer Nanomedicine Design and How to Make It Right. ACS Nano 14, 12281–12290 (2020). 10.1021/acsnano.9b09713

37 Pietras, E. M., et al. Functionally Distinct Subsets of Lineage-Biased Multipotent Progenitors Control Blood Production in Normal and Regenerative Conditions. Cell Stem Cell 17, 35–46 (2015). 10.1016/j.stem.2015.05.003

38 Singhal, S., et al. Origin and Role of a Subset of Tumor-Associated Neutrophils with Antigen-Presenting Cell Features in Early-Stage Human Lung Cancer. Cancer Cell 30, 120–135 (2016). 10.1016/j.ccell.2016.06.001

39 Saha, S. & Biswas, S. K. Tumor-Associated Neutrophils Show Phenotypic and Functional Divergence in Human Lung Cancer. Cancer Cell 30, 11–13 (2016). 10.1016/j.ccell.2016.06.016

40 Liu, C. Y., et al. Population alterations of L-arginase- and inducible nitric oxide synthase-expressed CD11b+/CD14(-)/CD15+/CD33+ myeloid-derived suppressor cells and CD8+ T lymphocytes in patients with advanced-stage non-small cell lung cancer. J Cancer Res Clin Oncol 136, 35–45 (2010). 10.1007/s00432-009-0634-0

41 Evrard, M., et al. Developmental Analysis of Bone Marrow Neutrophils Reveals Populations Specialized in Expansion, Trafficking, and Effector Functions. Immunity 48, 364–379 e368 (2018). 10.1016/j.immuni.2018.02.002

42 Muendlein, H. I., et al. Neutrophils and macrophages drive TNF-induced lethality via TRIF/CD14-mediated responses. Science Immunology 7 (2022). https://doi.org:ARTN eadd066510.1126/sciimmunol.add0665

43 Hackert, N. S., et al. Human and mouse neutrophils share core transcriptional programs in both homeostatic and inflamed contexts. Nature Communications 14 (2023). https://doi.org:ARTN 813310.1038/s41467-023-43573-9

44 Sharygin, D., Koniaris, L. G., Wells, C., Zimmers, T. A. & Hamidi, T. Role of CD14 in human disease. Immunology 169, 260–270 (2023). 10.1111/imm.13634

45 Pugin, J., et al. Cd14 Is a Pattern-Recognition Receptor. Immunity 1, 509–516 (1994). https://doi.org:Doi 10.1016/1074-7613(94)90093-0

46 Ding, Z. M., et al. Relative contribution of LFA-1 and Mac-1 to neutrophil adhesion and migration. J Immunol 163, 5029–5038 (1999).

47 Carlos, T. M. & Harlan, J. M. Leukocyte-Endothelial Adhesion Molecules. Blood 84, 2068–2101 (1994).

48 Fridlender, Z. G., et al. Polarization of Tumor-Associated Neutrophil Phenotype by TGF-β: “N1” versus “N2” TAN. Cancer Cell 16, 183–194 (2009). 10.1016/j.ccr.2009.06.017

49 Piccard, H., Muschel, R. J. & Opdenakker, G. On the dual roles and polarized phenotypes of neutrophils in tumor development and progression. Crit Rev Oncol Hematol 82, 296–309 (2012). 10.1016/j.critrevonc.2011.06.004

50 Zhang, J., et al. Engineering and Targeting Neutrophils for Cancer Therapy. Adv Mater 36, e2310318 (2024). 10.1002/adma.202310318

51 Kawai, T. & Akira, S. Toll-like Receptors and Their Crosstalk with Other Innate Receptors in Infection and Immunity. Immunity 34, 637–650 (2011). 10.1016/j.immuni.2011.05.006

52 Kawai, T. & Akira, S. The roles of TLRs, RLRs and NLRs in pathogen recognition. Int Immunol 21, 317–337 (2009). 10.1093/intimm/dxp017

53 Wicherska-Pawlowska, K., Wrobel, T. & Rybka, J. Toll-Like Receptors (TLRs), NOD-Like Receptors (NLRs), and RIG-I-Like Receptors (RLRs) in Innate Immunity. TLRs, NLRs, and RLRs Ligands as Immunotherapeutic Agents for Hematopoietic Diseases. Int J Mol Sci 22 (2021). https://doi.org:ARTN 1339710.3390/ijms222413397

54 Yan, H. M., Kamiya, T., Suabjakyong, P. & Tsuji, N. M. Targeting C-type lectin receptors for cancer immunity. Front Immunol 6 (2015). https://doi.org:ARTN 40810.3389/fimmu.2015.00408

55 Li, J. Y., et al. Tumor Cell-Intrinsic Factors Underlie Heterogeneity of Immune Cell Infiltration and Response to Immunotherapy. Immunity 49, 178-+ (2018). 10.1016/j.immuni.2018.06.006

56 Reynaud, D., et al. IL-6 Controls Leukemic Multipotent Progenitor Cell Fate and Contributes to Chronic Myelogenous Leukemia Development. Cancer Cell 20, 661–673 (2011). 10.1016/j.ccr.2011.10.012

57 Zhao, J. L., et al. Conversion of Danger Signals into Cytokine Signals by Hematopoietic Stem and Progenitor Cells for Regulation of Stress-Induced Hematopoiesis. Cell Stem Cell 14, 445–459 (2014). 10.1016/j.stem.2014.01.007

58 Tie, R. X., et al. Interleukin-6 signaling regulates hematopoietic stem cell emergence. Exp Mol Med 51 (2019). https://doi.org:ARTN 12410.1038/s12276-019-0320-5

59 Kimmel, B. R., et al. Potentiating cancer immunotherapies with modular albumin-hitchhiking nanobody-STING agonist conjugates. Nat Biomed Eng (2025). 10.1038/s41551-025-01400-0

60 Nagata, M., et al. A critical role of STING-triggered tumor-migrating neutrophils for anti-tumor effect of intratumoral cGAMP treatment. Cancer Immunol Immun 70, 2301–2312 (2021). 10.1007/s00262-021-02864-0

61 Xu, T. X., et al. Systemic administration of STING agonist promotes myeloid cells maturation and antitumor immunity through regulating hematopoietic stem and progenitor cell fate. Cancer Immunol Immun (2023). 10.1007/s00262-023-03502-7

62 Zanoni, I., Tan, Y. H., Di Gioia, M., Springstead, J. R. & Kagan, J. C. By Capturing Inflammatory Lipids Released from Dying Cells, the Receptor CD14 Induces Inflammasome-Dependent Phagocyte Hyperactivation. Immunity 47, 697-+ (2017). 10.1016/j.immuni.2017.09.010

63 Schröder, N. W. J., et al. Lipoteichoic acid (LTA) of and activates immune cells via toll-like receptor (TLR)-2, lipopolysaccharide-binding protein (LBP), and CD14, whereas TLR-4 and MD-2 are not involved. J Biol Chem 278, 15587–15594 (2003). 10.1074/jbc.M212829200

64 Roedig, H., et al. Biglycan is a new high-affinity ligand for CD14 in macrophages. Matrix Biol 77, 4–22 (2019). 10.1016/j.matbio.2018.05.006

65 Woodfin, A., et al. ICAM-1-expressing neutrophils exhibit enhanced effector functions in murine models of endotoxemia. Blood 127, 898–907 (2016). 10.1182/blood-2015-08-664995

66 Lee, S. H., et al. Intracellular Adhesion Molecule-1 Improves Responsiveness to Immune Checkpoint Inhibitor by Activating CD8 T Cells. Adv Sci 10 (2023). 10.1002/advs.202204378

67 Ma, V. P. Y., et al. The magnitude of LFA-1/ICAM-1 forces fine-tune TCR-triggered T cell activation. Sci Adv 8 (2022). https://doi.org:ARTN eabg448510.1126/sciadv.abg4485

68 Deeths, M. J. & Mescher, M. F. ICAM-1 and B7-1 provide similar but distinct costimulation for CD8+ T cells, while CD4+ T cells are poorly costimulated by ICAM-1. Eur J Immunol 29, 45–53 (1999).

69 Cox, M. A., Barnum, S. R., Bullard, D. C. & Zajac, A. J. ICAM-1-dependent tuning of memory CD8 T-cell responses following acute infection. P Natl Acad Sci USA 110, 1416–1421 (2013). 10.1073/pnas.1213480110

70 Eruslanov, E. B., et al. Tumor-associated neutrophils stimulate T cell responses in early-stage human lung cancer. Journal of Clinical Investigation 124, 5466–5480 (2014). 10.1172/Jci77053

71 Hines, J. B., Kacew, A. J. & Sweis, R. F. The Development of STING Agonists and Emerging Results as a Cancer Immunotherapy. Curr Oncol Rep 25, 189–199 (2023). 10.1007/s11912-023-01361-0

72 Motedayen Aval, L., Pease, J. E., Sharma, R. & Pinato, D. J. Challenges and Opportunities in the Clinical Development of STING Agonists for Cancer Immunotherapy. J Clin Med 9 (2020). 10.3390/jcm9103323

73 Wang, B., et al. Clinical applications of STING agonists in cancer immunotherapy: current progress and future prospects. Front Immunol 15, 1485546 (2024). 10.3389/fimmu.2024.1485546

74 Ramanjulu, J. M., et al. Design of amidobenzimidazole STING receptor agonists with systemic activity. Nature 564, 439–443 (2018). 10.1038/s41586-018-0705-y

75 Chin, E. N., et al. Antitumor activity of a systemic STING-activating non-nucleotide cGAMP mimetic. Science 369, 993–999 (2020). 10.1126/science.abb4255

76 Pan, B. S., et al. An orally available non-nucleotide STING agonist with antitumor activity. Science 369, 935-+ (2020). https://doi.org:ARTN aba609810.1126/science.aba6098

77 Zhao, H. Y., et al. Design of an Oral STING Agonist through Intramolecular Hydrogen Bond Ring Mimicking to Achieve Complete Tumor Regression. J Med Chem 68, 11365–11385 (2025). 10.1021/acs.jmedchem.5c00296

